# Diverse intrinsic properties shape transcript stability and stabilization in *Mycolicibacterium smegmatis*

**DOI:** 10.1101/2024.06.02.596988

**Authors:** Huaming Sun, Diego A. Vargas-Blanco, Ying Zhou, Catherine S. Masiello, Jessica M. Kelly, Justin K. Moy, Dmitry Korkin, Scarlet S. Shell

**Affiliations:** Program in Bioinformatics and Computational Biology, Worcester Polytechnic Institute, Worcester, Massachusetts, 01609, USA; Department of Biology and Biotechnology, Worcester Polytechnic Institute, Worcester, Massachusetts, 01609, USA

## Abstract

In mycobacteria, regulation of transcript degradation is known to occur in response to environmental stress and to facilitate adaptation. However, the mechanisms underlying this regulation are unknown. Here we sought to gain an understanding of the mechanisms controlling mRNA stability by investigating the transcript properties associated with variance in transcript stability and stress-induced transcript stabilization. We performed transcriptome-wide mRNA degradation profiling of *Mycolicibacterium smegmatis* in both log phase growth and hypoxia-induced growth arrest. The transcriptome was globally stabilized in response to hypoxia, with all transcripts having longer half-lives, but some having greater degrees of stabilization than others. The transcripts of essential genes were generally stabilized more than those of non-essential genes. We then developed machine learning models that utilized a compendium of transcript properties and enabled us to identify the non-linear collective effect of diverse properties on transcript stability and stabilization. The comparisons of these properties confirmed the association of 5’ UTRs with transcript stability, along with other differences between leadered and leaderless transcripts. Our analysis highlighted the protective effect of translation in log phase but not in hypoxia-induced growth arrest. Steady-state transcript abundance had a weak negative association with transcript half-life that was stronger in hypoxia, while coding sequence length showed an unexpected correlation with half-life in hypoxia only. In summary, we found that transcript properties are differentially associated with transcript stability depending on both the transcript type and the growth condition. Our results reveal the complex interplay between transcript features and microenvironment that shapes transcript stability in mycobacteria.

## INTRODUCTION

Regulation of mRNA degradation serves as a response mechanism of mycobacteria to energy-limited microenvironments. The mycobacteria include *Mycobacterium tuberculosis*, the causative agent of tuberculosis which led to over 1 million deaths in 2022 ^1^. Transcriptome-wide profiling of mRNA degradation in *M. tuberculosis* showed variance in transcript stability among genes and a global stabilization of the transcriptome in hypoxia, a stress condition that *M. tuberculosis* encounters within granulomas during infection ^2–4^. However, the regulatory mechanisms that govern transcript stability and stress-induced stabilization remain poorly understood. Further study of the regulation of mRNA degradation in mycobacteria is needed to facilitate our understanding of the stress response strategies that *M. tuberculosis* employs to adapt and persist within the host.

Global mapping of transcription start sites revealed that approximately 25% of RNA transcripts lack a 5’ untranslated region (5’ UTR) in mycobacteria (referred to as leaderless transcripts) ^5–7^. Studies have shown that the presence of Shine-Dalgarno (SD) ribosome binding site sequences within the 5’ UTR is associated with higher mRNA expression levels in bacteria, measured by RNAseq expression data ^5,7^. However, it is also known that some leaderless transcripts have comparable translation efficiencies with leadered transcripts in both *M. smegmatis* and *M. tuberculosis* ^6,8^, potentially due to an alternative translation initiation mechanism and unique RNA characteristics, such as less structured start codon regions ^9,10^. In *M. tuberculosis*, SD-independent translation of leaderless transcripts is less affected during adaptation to stress environments than canonical leadered translation ^5,11^. While it is likely that the 5’ regions of transcripts can contribute to the variability in mRNA half-life through either translation efficiency or degradation initiation ^5,7,12–18^, the mechanisms are not fully characterized.

Various transcript properties (features) have been reported to be associated with mRNA stability either transcriptome-wide or for individual transcripts in various organisms. Transcriptome-wide associations with mRNA stability have been shown for growth rate in *L. lactis* and *E. coli* ^19,20^, transcript abundance in *E. coli* and *L. lactis* ^21,22^, GC content in *B. cereus*, *E. coli* and *S. cerevisiae* ^21,23,24^, 5’ UTR-related features in *S. cerevisiae* and *E. coli* ^21,25^, 3’ UTR-related features in *S. cerevisiae* ^25^, gene function and essentiality in *B. cereus* and *E. coli* ^21,23^, transcript length in *L. lactis*, *E. coli*, and *S. cerevisiae* ^19,21,24^, ribosome density in *S. cerevisiae* ^24^, and adjacent codon pair usage in *S. cerevisiae* ^26^. The impacts of some transcript features such as start codon identity and GC content in *S. cerevisiae* ^24^, 5’ UTR-related features in *L. lactis*, *A. baumannii*, *H. pylori*, *E. coli*, *B. subtilis*, and *M. smegmatis* ^8,12,22,27–35^, and transcript abundance in *E. coli* and *L. lactis* ^22^ were also validated experimentally with individual transcripts. In mycobacteria, the transcript features that impact mRNA degradation rates are largely unexplored, with existing analysis limited mainly to a few transcript features and their individual broad correlations with transcript half-life in log-phase growing *M. tuberculosis* ^2^. It is unknown which of the wide range of potentially associated features impact transcript stability, how the impacts of these features interact, and how they differ according to growth condition.

To model the underlying collective effect of multiple transcript features on stability, recent studies in *E. coli* and *S. cerevisiae* have used linear regression models to incorporate multiple features and quantify their contributions to variance in degradation rates ^21,24,25^. However, a limitation is that these models simplify the relationship between the features by assuming that they can be combined linearly to determine transcript half-life. Although more advanced sequence-based machine learning models could also be applied to predict stability, their performances rely on large amounts of data for training with the focus more on achieving accurate prediction rather than understanding the underlying mechanisms ^36,37^.

Given our lack of understanding of the impact of RNA features on mRNA degradation in mycobacteria, we sought to develop a comprehensive machine learning framework to identify the transcript properties associated with transcript half-life in the model organism *Mycolicibacterium* (nee *Mycobacterium*) *smegmatis* in both log phase growth and hypoxia-induced growth arrest. We found that in contrast to some previous reports on *M. tuberculosis* and *E. coli*, no single feature had a dominant association with mRNA half-life; rather, half-lives were best explained by the non-linear interactions of many features. The features that best explained transcript half-lives differed between log phase growth and hypoxia, and while the half-lives of most transcripts were longer in hypoxia, those of essential genes were lengthened the most. Features associated with efficient translation were generally predictive of longer half-lives in log phase but not in hypoxia, consistent with the idea that translation protects mRNA from degradation in rapidly growing cells and lower levels of translation in non-growing cells limit its impact. mRNA secondary structure was also generally predictive of longer half-lives in cases where it did not negatively impact translation, but in ways that varied by condition and transcript leader type. 5’ UTR features were predictive of half-life in ways that appeared to extend beyond mediating translation initiation. Surprisingly, gene length was predictive of slower degradation in hypoxia, consistent with models in which diffusion of large molecules is slower in non-growing cells. Taken together, our results reveal the landscape of the collective effect of diverse transcript features on stability under different conditions in *M. smegmatis*.

## MATERIAL AND METHODS

### Strains and culture conditions to generate transcriptome-wide mRNA degradation datasets

Transcriptomic data were obtained from *M. smegmatis* strain SS-M_0424, a derivative of mc^2^-155 previously described ^38^, in which a *hyg^R^* gene was inserted upstream of, and divergent from, the *rne* gene promoter, and a *kan^R^*-marked plasmid expressing tetR38 was integrated at the L5 site. This strain was constructed as a control for an *rne* knockdown strain, and its genetic modifications did not affect expression of *rne*. *M. smegmatis* was grown at 37° C with 200 rpm shaking in Middlebrook 7H9 broth supplemented with final concentrations of 0.2% glycerol, 0.05% Tween-80, 3 mg/L catalase, 2 g/L glucose, 5 g/L bovine serum albumen fraction V, and 0.85 g/L sodium chloride. RNA was extracted from cultures at defined time-points following addition of 150 µg/mL rifampicin to block transcription initiation. The log phase cultures are described in ^38^. Cultures for hypoxia were sealed in vials as described in ^39^. The volume of culture in each bottle was 13.5 mL and the OD at the time of sealing the bottles was 0.01. 19 hours after sealing the bottles, rifampicin was injected through the rubber cap with a needle, and at the indicated timepoints (0, 3, 6, 9, 15, 30 and 60 minutes) bottles were opened, contents poured into 15 mL conical tubes, and the tubes submerged in liquid nitrogen. The elapsed time between opening the hypoxia bottles and submerging the cultures in liquid nitrogen was approximately 6 seconds. Frozen cultures were stored at -80° C. Cultures were thawed on ice and RNA extracted as in ^39^. The RNAs were submitted to the Broad Institute Microbial ‘Omics Core where Illumina libraries were constructed as described in ^40^ and sequenced. The hypoxia cultures were grown and their RNA extracted and sequenced together with the log phase cultures described in ^38^. Separately, the RNA samples were used to synthesize cDNA and perform quantitative PCR as described in ^39^ and the resulting data were used for normalization of the RNAseq data as described in ^38^.

### *M. smegmatis* genome sequence and gene annotations

The transcript features were quantified using the genome sequence of *M. smegmatis* strain mc^2^-155 (NC_008596.1) from Mycobrowser Release 4 ^41^. The gene annotations were updated as described ^38^ and listed in Table S1. Using these annotations, we defined 1939 leadered transcripts (Table S1) and 960 leaderless transcripts (Table S1) with high confidence. For transcripts with multiple transcription start sites (TSSs), the leadered type was determined by the TSS with the highest read coverage in log phase ^7^.

### RNAseq data processing and half-life calculations

RNAseq data were processed and normalized to produce transcript degradation profiles for log phase and hypoxia as described ^38^. Log phase half-lives were calculated as described ^38^. To calculate half-lives in hypoxia with high-confidence, we only used genes for which linear regression of log2 transcript abundance between first 5 time points (0, 3, 6, 9, 15 mins) had a mean squared error (MSE) < 0.5 ^38^. Genes with zero read counts for any replicate for any of those 5 time points were also excluded. Half-life was calculated as -1/slope.

### UMAP visualization of transcript degradation profiles in log phase and hypoxia

The log2 normalized RNAseq coverage of 7120 genes in 42 samples (3 replicates of each of the 7 time points in log phase and hypoxia) were used to visualize the transcript degradation profiles in *M. smegmatis*. UMAP plots were made in `R v4.3.2` using package `umap v0.2.10.0` with the parameters `n_neighbors = 20, min_dist = 0.25, n_component = 2, random_state = 77`. The seed value in R was set to be 7.

### Transcript property quantification

To identify the transcript properties that are associated with degradation, we quantified many potential candidate properties (Tables S2 and S3) for each of the 7120 CDSs (Table S1).

*Nucleotide frequency.* This group of properties was quantified through the nucleotide frequency percentage relative to 5’ UTR or CDS region length. It contains usage of single nucleotides, adjacent dinucleotide motifs, and total G+C content. For each CDS, we quantified nt frequency for the 5’ 18 nt, the 3’ 18 nt, and the entire CDS as separate properties.

*Codon frequency.* In addition to the percentage of each nonstop codon calculated in the same manner as the nucleotide frequency, we added binary indicators for the choice of start codon (AUG, GUG and UUG) and stop codon (UAA, UAG and UGA). We calculated the Codon Pair Bias (CPB) using the Codon Pair Score (CPS) of all codon pairs that make up the CDS ^42^. The CPS was calculated for each of the 3904 possible codon pairs (61 * 64), including stop codons only being used as the second codon to capture potential bias at the 3’ end of the CDS.

RNA *Secondary structure.* These properties were quantified using the `ViennaRNA v2.5.0` package ^43^ through the following three metrics: the ΔG of the minimum free energy (MFE) structure for a given transcript segment, the number of unpaired nucleotides at the 5’ end of the MFE structure, and the probability of specific nucleotides near the 5’ end being unpaired. The ΔGs of MFE structures were calculated using `RNAfold v2.5.0`. To overcome the positive correlation between ΔG and sequence length, we calculated ΔG of MFE (ΔGMFE) structures in a sliding window manner. Each sequence was split into subsequences by *M* nt windows, each with *M/2* nt overlap. The ΔGMFE for a given sequence is the averaged ΔGMFE of all its subsequences generated by a sliding window. For 5’ UTRs and CDSs, we divided each sequence into thirds and used such sliding window ΔGMFEs to quantify the predicted structure of the 5’ third, middle third, and 3’ third as well as the entire sequence. For the 5’ UTRs, we sought to distinguish between secondary structure directly affecting ribosome binding and other secondary structure. We therefore excluded the 3’-most 15 nt before dividing the sequence into thirds. Additionally, only 5’ UTR sequences longer than 35 nt before removing the 15 nt ribosome binding site were used, and only a 20 nt sequence window was used. For the CDS region, we calculated ΔGMFEs using 20, 50, and 100 nt windows. The 3’ UTRs were approximated as 60 nt after the stop codons. The MFE for 3’ UTRs were calculated using a 20 nt window only.

We also used the ΔGMFE structures to measure the accessibility of the mRNA translation initiation region (TIR) for ribosome binding ^10^. We calculated Δ*Gunfold* separately for leadered transcripts that have 5’ UTRs at least 12 nt long (1809 transcripts) and leaderless transcripts (960 transcripts). To calculate Δ*Gunfold*, Δ*GmRNA* was first calculated using `RNAfold v2.5.0` to represent the folded state of the mRNA TIR in the absence of ribosome binding. For leadered transcripts with 5’ UTRs at least 25 nt long, this region was defined as 25 nt upstream of the start codon and the first 25 nt of the coding sequence. For leadered transcripts with 5’ UTRs shorter than 25 nt and for the leaderless transcripts, this region was defined as 50 nt downstream of the transcription start sites (TSSs). Then to approximate the ribosome-bound state of the mRNA TIR, Δ*Ginit*, the TIR structure prediction was processed using `RNAstructure v6.3` to break any base pairing within the ribosome footprint. The ribosome footprint was assumed to be 12 nt upstream of the start codon and the first 13 nt of the CDS for leadered transcripts, and the first 13 nt of the CDS for leaderless transcripts ^44^. Then Δ*Gunfold* was calculated as Δ*Ginit* - Δ*GmRNA*.

The number of unpaired nucleotides at each transcript 5’ end was predicted from MFE structures produced by `RNAfold v2.5.0` when folding the entire CDS (leadered and leaderless genes), the 5’ UTR (leadered genes only), the 5’ UTR plus the first 18 nt of the CDS (leadered genes only), or the first 20 nt of the transcripts (leadered and leaderless genes).

Separately, the probabilities of certain transcript regions being unpaired were predicted using `RNAplfold v2.5.0.` To assess the base-pairing status of the 5’ ends of transcripts in a different way, we folded the first 20 nt of each transcript and calculated for the first 3 nt and 5 nt (i) the probabilities that all the nucleotides are unpaired or (ii) the averaged unpaired probability of each nucleotide. To predict accessibility of ribosome-binding regions in leadered transcripts, we folded the last 30 nt of the 5’ UTR plus either the first 20 nt of the CDS or the start codon only. We then quantified the probability of the entire start codon being unpaired as well as the averaged dinucleotide unpaired probability over either the entire folded sequence or the Shine-Dalgarno region (-6 to -14 relative to the start codon) ^6^.

*Ribosome occupancy.* We used RNAseq data from GSE127827, which included libraries made from total rRNA-depleted RNA (referred to henceforth as mRNA libraries) and well as libraries made from ribosome footprints. After retrieving data using `SRA Toolkit v3.0.0`, we processed the sequencing data following the original methods with some modifications ^45^. First, quality control was performed using `FastQC v0.11.9` ^46^. The ribosome footprint data were further processed using `Trimmomatic v0.39` with the options `ILLUMINACLIP:∼/adaptors_SE.fa:2:30:10 SLIDINGWINDOW:4:20 MINLEN:25`, which including removing adaptors, cutting reads when average quality per nucleotide was lower than 20 within a 4-nt sliding window, and discarding reads less than 25 nt long ^47^. This was not necessary for mRNA libraries due to their higher quality. Next, for both ribosome footprint and mRNA libraries, we performed alignment to discard reads aligned to tRNA and rRNA using `Bowtie2 v2.4.5` with the option `--very-sensitive` ^48^. The remaining reads were then aligned to the genome sequence of *Mycolicibacterium smegmatis* strain mc^2^-155 using `Bowtie2 v2.4.5` with the option `--sensitive-local`. Reads and alignments were processed and sorted using `SAMtools v1.16.1` ^49^. To further remove unmapped reads, PCR or optical duplicate reads, reads that are not primary alignments and alignments with MAPQ smaller than 10, we filtered the alignments using `SAMtools v1.16.1` with the options `-q 10 -F 1284`. The remaining alignments were then quantified in TPM for both ribosome footprint and mRNA libraries using `StringTie v2.2.1` ^50^. Such quantification was done separately for four different transcript regions: (i) the entire CDS, (ii) the entire CDS plus the 20 nt upstream, (iii) the 5’ end of the transcript (the first 18 nt of the CDS for leaderless transcripts or the last 20 nt of the 5’ UTR plus the first 18 nt of the CDS for leadered transcripts), and (iv) the CDS excluding its first 18 nt. We also quantified the coverage for each third of every CDS (5’ third, middle third, 3’ third) to capture regional differences. Then for each of the two affinity tag swapped strains, we calculated the TPM ratios of ribosome footprint library coverage over mRNA library coverage using the averaged TPMs of two replicates. At the end, the normalized ribosome occupancy for each CDS in these transcript regions was calculated as the averaged TPM ratios of these two dual-RpsR-tagged strains.

*Shine-Dalgarno sequence*. To approximate the strength of the Shine-Dalgarno sequences of leadered genes, we quantified the GA percent and GA frequency within the region of -17 to -4 relative to the start codon, as well as the frequencies of 17 specific Shine-Dalgarno motif variants in the 25 nt upstream of the start codon. GA percent is the percentage of Gs and As in the sequence. GA count is the total frequency of all di-nucleotide sequences (GG, AA, AG, and GA) in two “reading frames” of the sequence, one starting at the first nt and one starting at the second nt of the sequence.

*Other properties.* This group of properties includes the sequence length and steady-state transcript abundance (0 min RIF treatment; “initial abundance”). We quantified length for 5’ UTRs and CDSs based on the annotation of 7120 CDSs (Table S1). CDS abundance was normalized by CDS length. The initial abundances in log phase and hypoxia were used respectively for log phase and hypoxia model development.

### Feature selection procedure

Our complete feature set includes five different feature types (nucleotide frequency, codon frequency, secondary structure, ribosome occupancy, and others) in four transcript regions (5’ UTR, 5’ end of transcript, CDS, and 3’ UTR) (Figure 3A; Table S2 and S3). The design of this feature set was driven by our hypothesis that the transcript stability is controlled by the unknown combination of multiple transcript properties. However, the intersection of multiple feature types within and across transcript regions leads to high correlations among several features. Although those correlations might not directly affect machine learning model performance, the shared credit of correlated features contributing to the predictions could affect the importance rankings of features. Such correlations also complicate the interpretation of feature contributions.

We therefore sought a feature selection algorithm to minimize the influence of the correlated features without losing potentially important features. Commonly used selection techniques lack the ability to consider both the relationships among the features themselves and the relationships between the features and the predicted class ^51^. To perform feature selection in a manner suitable for our feature structure and goals, we developed the following algorithm. Our algorithm only targeted the highly correlated features (| Spearman’s *ρ* |>= 0.6) that were of the same type and within the same transcript region. We also took into account correlations between the individual features’ values and classes, measured by the Kendall rank correlation coefficient ^52^. For this process, we considered the 5’ UTR and 5’ end of the transcript to be the same transcript region. Our goal was to select features that have less correlations with other features, while potentially contributing the most to the model performance. The selection procedure was done separately for each of six models that used the combined feature set to predict half-life class: leadered genes and leaderless genes each in log phase, hypoxia, and fold change in hypoxia relative to log phase. The algorithm returned a list of features to be used for the machine learning model training and evaluation (See Supplementary Materials).

### Machine learning classifier training

Given the limited number of leadered and leaderless transcripts and the imbalanced number of genes in the half-life classes, random forest emerged as an optimal choice given its fast training convergence and ability to avoid overfitting ^53^. To train and evaluate the classifiers, we implemented 5-fold nested cross-validation using the `scikit-learn 1.2.1` package ^54^.

Each dataset was split into 5 folds in a stratified manner for outer cross-validation. The same training and testing sets were used for random class prediction models and random forest classifiers at each fold iteration to compare their performances. To measure the performance of the classifiers given the numerically imbalanced yet equally important classes, we used the macro F1 score, *i.e.*, the unweighted mean of F1 scores for each class. See below formula for the F1 score and the macro F1 score.

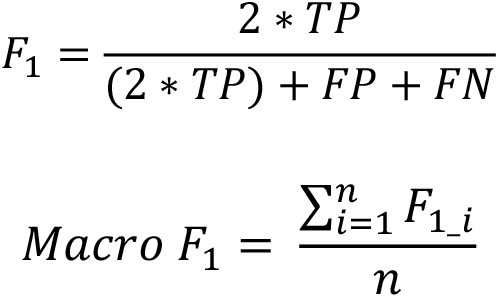

Where *TP* is the number of true positives, *FN* is the number of false negatives, and *FP* is the number of false positives. *n* is the number of classes.

In order to perform hyperparameter tuning for random forest classifiers, the training set was split in the same manner as before for inner cross-validation to do a randomized search on hyperparameter sets ***H*** (max_depth: [3, 5, 7], min_samples_leaf: [20, 30, 50], min_samples_split: [5, 10, 20]). The optimal hyperparameters set ***h\**** selected by inner cross-validation was used for training with the entire training set to obtain the optimal model ***RandomForestModel\****, which was then evaluated using the outer testing set. To quantify the contributions of individual features, we used both the impurity-based Gini importance and the SHAP values of predictions made on outer testing sets. To reduce the bias of random sampling, the nested cross-validation was repeated 10 times with a different random split of training and testing sets each time. Ultimately, the outputs of the nested cross-validation include the F1 scores of all fold iterations for 10 repetitions; the averaged F1 scores of the 10 repetitions; averaged Gini importance scores of 10 repetitions; and SHAP values for each stability class across all fold iterations for 10 repetitions (See Supplementary Materials).

### Statistical comparison of machine learning models

For machine learning models developed using cross-validation, there are two potential issues in testing the statistical significance of model performance differences. First, the performance of classifiers could be driven by a specific split of training and testing sets. To better ensure that a potential significant difference between classifier F1 scores is not due to a random split, ideally the difference in F1 score should be calculated using the same fold iteration of data for each pair of compared classifiers. Second, in the case of statistically testing the significance of differences between two distributions of F1 scores or F1 score differences, the commonly used Student’s t-test could provide misleading results in the context of cross-validation. The reason is that the resampled data in training and testing makes the F1 scores, and thereby the F1 score differences, dependent across iterations. This violates the independence assumption in the Student’s t-test, and could lead to high Type I error due to the underestimated variance of difference ^55^. To address these problems, Nadeau and Bengio proposed a corrected paired t-test, which can take into account the dependency in samples and reduce the number of fold positive errors ^56^. In our case, the classifiers were trained and evaluated using 5-fold nested cross-validation. The entire procedure was repeated 10 times to get averaged performance. Each time, the model was trained and tested using the subsampled training and testing sets that were overlapped in different fold iterations. We implemented Nadeau and Bengio’s corrected paired t-test to evaluate the differences between our random forest classifiers and random estimators that predict class membership randomly without using any of the features. Each classifier and a random estimator are trained and tested using the same fold of the dataset through the cross-validation. At the end, 50 pairs of F1 scores (5 folds * 10 repetitions) for these two classifiers were collected to test for the significance of difference. For the classifiers that were trained separately for the genes, conditions, or features being compared, we were not able to train and evaluate their performances with the same dataset. We therefore calculated ΔF1 score as the difference between F1 score from the random forest classifier and the random estimator for each comparison of interest and used the Wilcoxon rank-sum test to compare the ΔF1 scores from 10 repetitions between conditions, gene types, and features.

### Essential gene enrichment analysis

Essentiality of 6642 *M. smegmatis* genes were defined using the CRISPR interference system ^57^. To statistically test the enrichment of essential genes in each stability class, only genes with essentiality designations and half-lives calculated in both log phase and hypoxia were used.

This resulted in 3680 genes, of which 1327 were classified as leadered, and 793 were classified as leaderless. These genes were tested for essentiality enrichment in half-life classes using a hypergeometric test with FDR correction for multiple hypothesis testing (Figure 2G).

## RESULTS

### Overview of an experimental and computational framework to unravel the intrinsic transcript properties that impact transcript stability in *M. smegmatis*

Bacterial mRNA half-lives are known to vary among transcripts and between conditions. To identify the transcript properties that contribute to the variance in transcript stability within and across conditions in *M. smegmatis*, we developed an experimental and computational framework consisting of the following four stages (Figure 1). We will summarize the stages here and describe them in greater detail in subsequent sections. First, we used RNA-seq to quantify transcript half-lives transcriptome-wide. To characterize the impact of the microenvironment on transcript stability, transcript degradation profiles were determined in log phase growth and hypoxia-induced growth arrest. We calculated transcript half-lives by linear regression of log2 transcript abundance over time for each condition. High-confidence half-lives were determined for 4,857 genes in log phase and 4,864 genes in hypoxia. The log phase half-lives were published previously in ^38^. Second, transcripts were classified into quartiles based on half-life in log phase or hypoxia, or by fold-change in half-life in hypoxia compared to log phase. Third, we compiled transcript properties (features) that we hypothesized could affect half-life and developed random forest classifiers to identify properties predictive of half-life class membership. This was done separately for leadered and leaderless genes given differences in their features and the idea that their half-life determinants might differ. Fourth, the values of features identified as important were plotted by half-life class to provide an overview of the association between transcript properties and classes. We also implemented SHAP (SHapley Additive exPlanations) during classifier development to further explore the impact of features on each class ^58^. Together, our pipeline reveals a comprehensive landscape of transcript half-lives and the transcript features influencing these half-lives in *M. smegmatis* in commonly studied rapid-growth and growth-arrested conditions.

**Figure 1.**
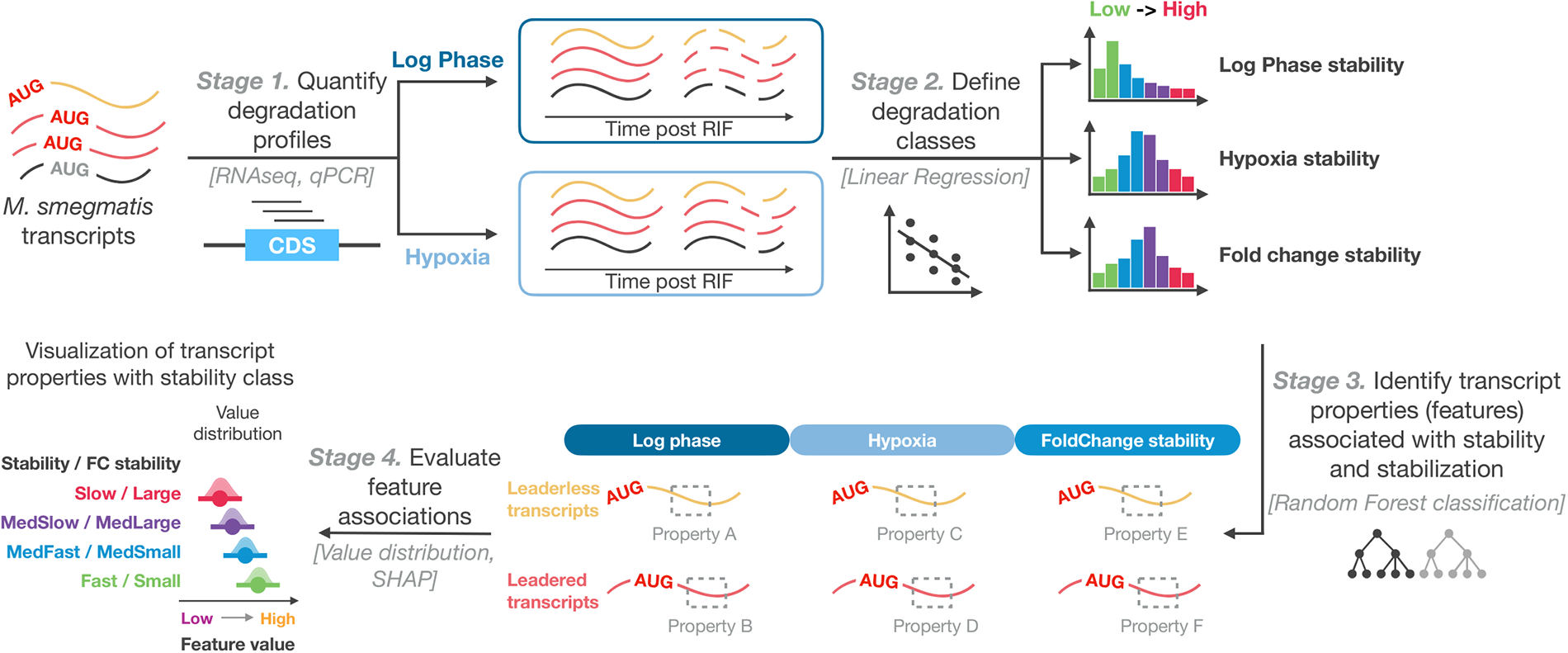
Schematic of the framework to identify transcript properties that impact transcript stability in *M. smegmatis*. The framework was designed to reveal the transcript properties that were differentially associated with transcript stability depending on the transcript type and condition. Stage 1: Transcriptome-wide mRNA degradation profiles were collected in log phase and hypoxia using RNAseq. Stage 2: In each condition, transcripts were classified into four groups according to their half-lives. Stages 3 and 4: A series of random forest classifiers were trained to classify transcripts into their assigned half-life class based on the values of a set of transcript properties (features), and identify the features important for these classifications.

### Transcript degradation profiles capture variance in transcript stability both within and between growth conditions

To obtain transcriptome-wide mRNA degradation profiles in *M. smegmatis*, we inhibited transcription initiation with rifampicin (RIF) followed by RNA extraction at various time-points. Hypoxia was produced by a variation of the Wayne model in which cultures were sealed with a defined volume of headspace and incubated with shaking for 19 hours ^39^. RIF was injected through rubber caps with a needle to minimize introduction of oxygen, and bottles were sacrificed at each time-point. Transcript abundance was quantified by RNAseq for samples harvested after 0, 1, 2, 4, 8, 16, and 32 minutes of RIF exposure in log phase and 0, 3, 6, 9, 15, 30, and 60 minutes of RIF exposure in hypoxia. Transcript abundance was normalized by relative abundance values determined for a set of genes by qPCR ^38^. A two-dimensional overview of the degradation profiles obtained by UMAP revealed a global difference between log phase and hypoxia (Figure 2A) ^59^, consistent with expectations from previous work indicating the hypoxia causes changes in gene expression patterns as well as longer transcript half-lives in mycobacteria ^2,7,39,60,61^. Samples also clustered by time-point after addition of RIF, corresponding with the temporal changes in transcript abundance and indicating that our method successfully captured the global degradation trends in both conditions.

**Figure 2.**
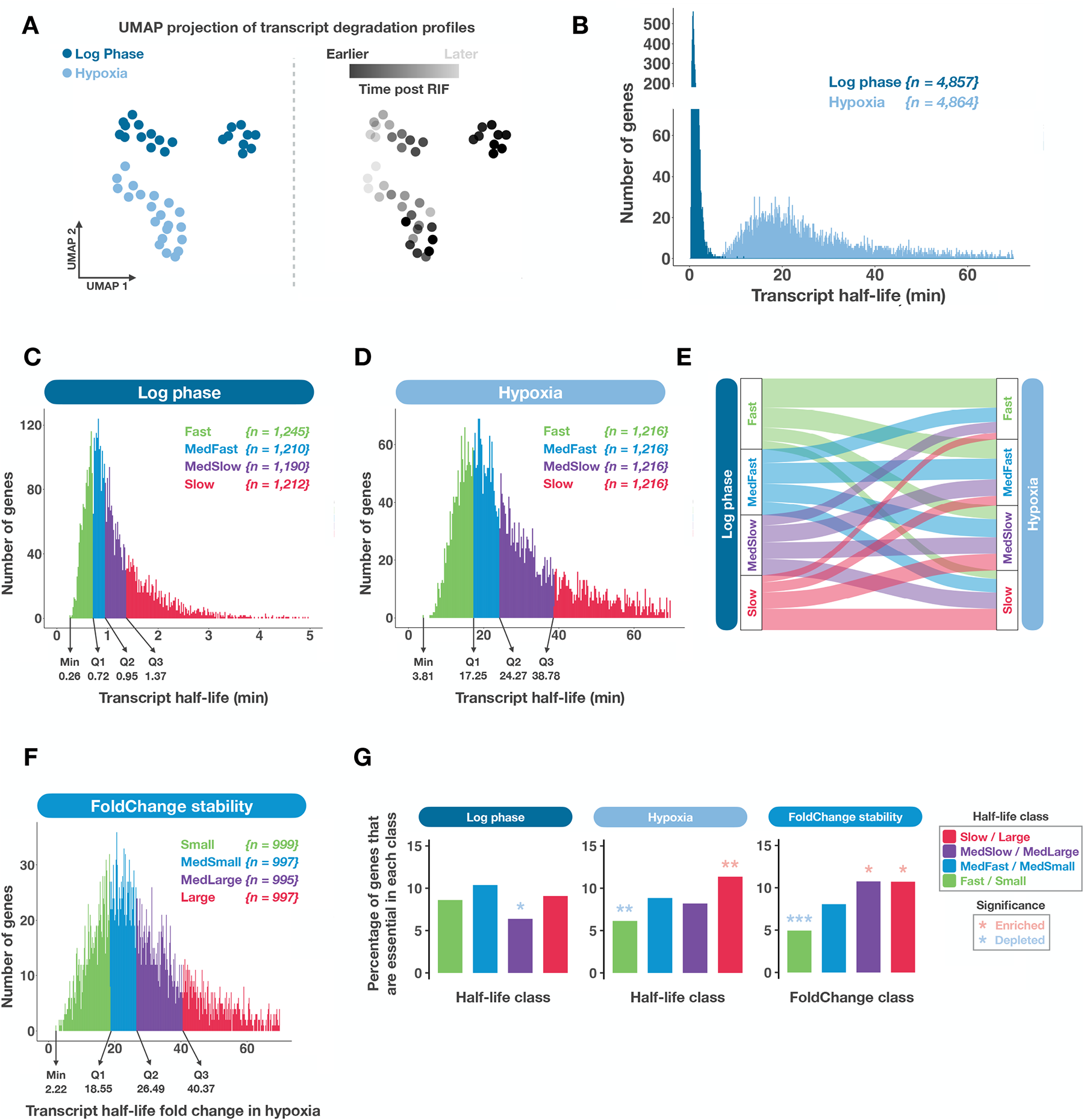
Transcriptome-wide mRNA degradation profiles in *M. smegmatis*. **A.** UMAP projection showing condition differences and temporal changes in global degradation profiles. Each dot represents an RNAseq library from which normalized mRNA abundance values for each gene were obtained. The same data are shown in the two UMAP panels, colored according to condition (left, two conditions) or time after adding RIF (right, time points). **B.** Distributions of transcript half-lives in log phase and hypoxia. Distribution plots were made in `R v4.3.2` using package `ggbreak v0.1.2` ^79^. **C-D.** Half-life distributions with classes defined by half-life quartiles in log phase and hypoxia. **E.** Comparison of half-life class membership between log phase and hypoxia. **F.** Distribution of half-life fold changes in stabilization with classes defined by fold change quartile. **G.** Frequency of essential genes in each half-life class. Significance of enrichment and depletion of essential genes within each class were tested using a hypergeometric test with FDR correction (Materials and Methods). *p.adjust* < 0.05 *, *p.adjust* < 0.01 **, *p.adjust* < 0.001 ***.

To further describe the transcript degradation process, we used linear regression models to calculate transcript half-lives from the degradation profiles (Figure 2B; Table S1). The time-points used for half-life determination were carefully chosen to avoid confounding from continued elongation by RNA polymerase following addition of RIF as well as decreases in degradation rate that appear to be induced by RIF over time ^38^. As expected, there were wide ranges of half-lives in each condition. The half-life measurements also confirmed the expected global difference between log phase and hypoxia, with stabilization of all transcripts evident in hypoxia (Figure 2B). These findings are consistent with a previous assessment of bulk transcript stability in hypoxia-exposed *M. tuberculosis* ^2^, and our previous work showing stabilization of several transcripts in hypoxia-exposed *M. smegmatis* ^39^. However, this is the first transcriptome-wide report of mRNA half-lives in any hypoxia-exposed mycobacterial species. The observed global variance in transcript stability and transcript stabilization in response to hypoxia was maintained when we examined the transcripts of subsets of genes with defined transcription start sites (TSSs) (Supplementary Figure S1A, C). These subsets were composed only of genes that were monocistronic or the first in a polycistron, according to the annotations in ^7^ (Materials and Methods), and classified as leadered (having a 5’ UTR of 5 nt or more) or leaderless (lacking a 5’ UTR). Genes that were second or beyond in a polycistron or lacked annotated TSSs were excluded. For genes with multiple TSSs, we used the TSS with the highest read coverage in log phase to define the 5’ UTR or lack thereof ^7^. Direct comparison of leadered and leaderless transcripts showed a statistically significant yet biologically limited difference in half-lives, with more leadered transcripts having longer half-lives in both log phase and hypoxia (Supplementary Figure S1E-H).

To facilitate construction of machine learning models to identify transcript features affecting half-life, we grouped transcripts into classes based on half-life. Since the classes were split from a continuous range of half-lives, in theory one could define any number of classes. To select the number of classes that most accurately represented the degradation landscape, we first performed hierarchical clustering of the degradation profiles (Materials and Methods). The clustering produced four major classes with distinct degradation patterns (Supplementary Figure S2A, E). We therefore decided to create four half-life classes. We chose to define classes based on half-life quartiles rather than by clustering of the complete degradation profiles to avoid confounding from continued elongation of RNA polymerase after addition of RIF as well as RIF-induced stress responses (Figure 2C, D). Nonetheless, the classes defined by half-lives had very similar gene composition to the clusters defined by hierarchical clustering (Supplementary Figure S3A-F) and similarly separated genes according to transcript degradation rate (Supplementary Figure S2).

When comparing the gene sets in each half-life class in log phase and hypoxia, we found that many genes switched classes in the two conditions (Figure 2E). This was true for both leadered and leaderless transcripts (Supplementary Figure S1B, D). This suggested that the relationship between transcript features and half-life differs in different conditions. To facilitate later identification of those features, we additionally classified genes according to the extent of stabilization in hypoxia vs log phase (defined by fold-change in half-life, Figure 2F).

Interestingly, we found that in hypoxia, genes classified as essential by CRISPR interference ^57^ were significantly enriched in the slowest degradation class while significantly depleted in the fastest degradation class (Figure 2G). Consistent with this, genes with a larger fold-change in stability in response to hypoxia were more likely to be essential than those with a smaller fold-change (Figure 2G). However, there was no consistent relationship between essentiality and half-life in log phase. This result supports the idea that global transcript stabilization in response to hypoxia is likely a regulatory mechanism as well as an energy-saving mechanism in mycobacteria. The significant stabilization of essential genes in hypoxia was only observed for leadered genes and not for leaderless genes, which suggests the possibility of different regulatory mechanisms for those two types of transcripts (Supplementary Figure S4A, B).

### Nonlinear combinations of transcript properties (features) appear to specify half-life

We sought to identify the transcript properties that specify transcript half-life in *M. smegmatis*. To address this question as agnostically as possible, we compiled and quantified hundreds of properties, which we refer to as features. These included nucleotide and sequence features, predicted secondary structure features, and other features such as length, steady-state abundance, and ribosome occupancy from a published dataset ^45^. We categorized the features by type as well as by the gene region under consideration (Figure 3A), because we expected that some features would have different impacts depending on their location; for example, A/U-rich codons are expected to promote translation when located near the start codon due to their impact on secondary structure ^62,63^, but be translated less efficiently when located elsewhere in a coding sequence due to being less preferred codons in mycobacteria.

Random forest classifiers were then trained separately for each transcript type (leadered and leaderless) in each condition (log phase, hypoxia, and fold change in hypoxia relative to log phase) (Figure 3B). The classifiers were trained using 5-fold nested cross-validation and evaluated by the difference in F1 score compared to random prediction models (⊗F-score; see Materials and Methods). We trained classifiers using combined feature sets as well as using only features of each of the six types (5’ UTR, CDS nucleotide, CDS secondary structure, codon, translation, and others) in order to evaluate the contribution of each feature type (Figure 3B). For the combined feature sets, we used a customized feature selection procedure to reduce the number of correlated features (Materials and Methods). As we predicted, classifiers that used the combined feature sets achieved the best performances, suggesting that the stability of transcripts is specified by the combination of various types of transcript properties. The ⊗F-scores were low compared to those typically reported for random forest classifiers designed to distinguish between distinct clinical or physiological states (*e.g.*, diseased tissue vs healthy tissue), but were consistent with expectations for our data type, in which classes were made from continuous distributions of half-life values. A majority of the classifiers performed significantly better than random, and were strong enough to facilitate our overarching goal of identifying the features that impact half-life. Interestingly, most of the feature types could individually predict transcript stability with performance that varied depending on transcript type and condition (Figure 3B).

**Figure 3.**
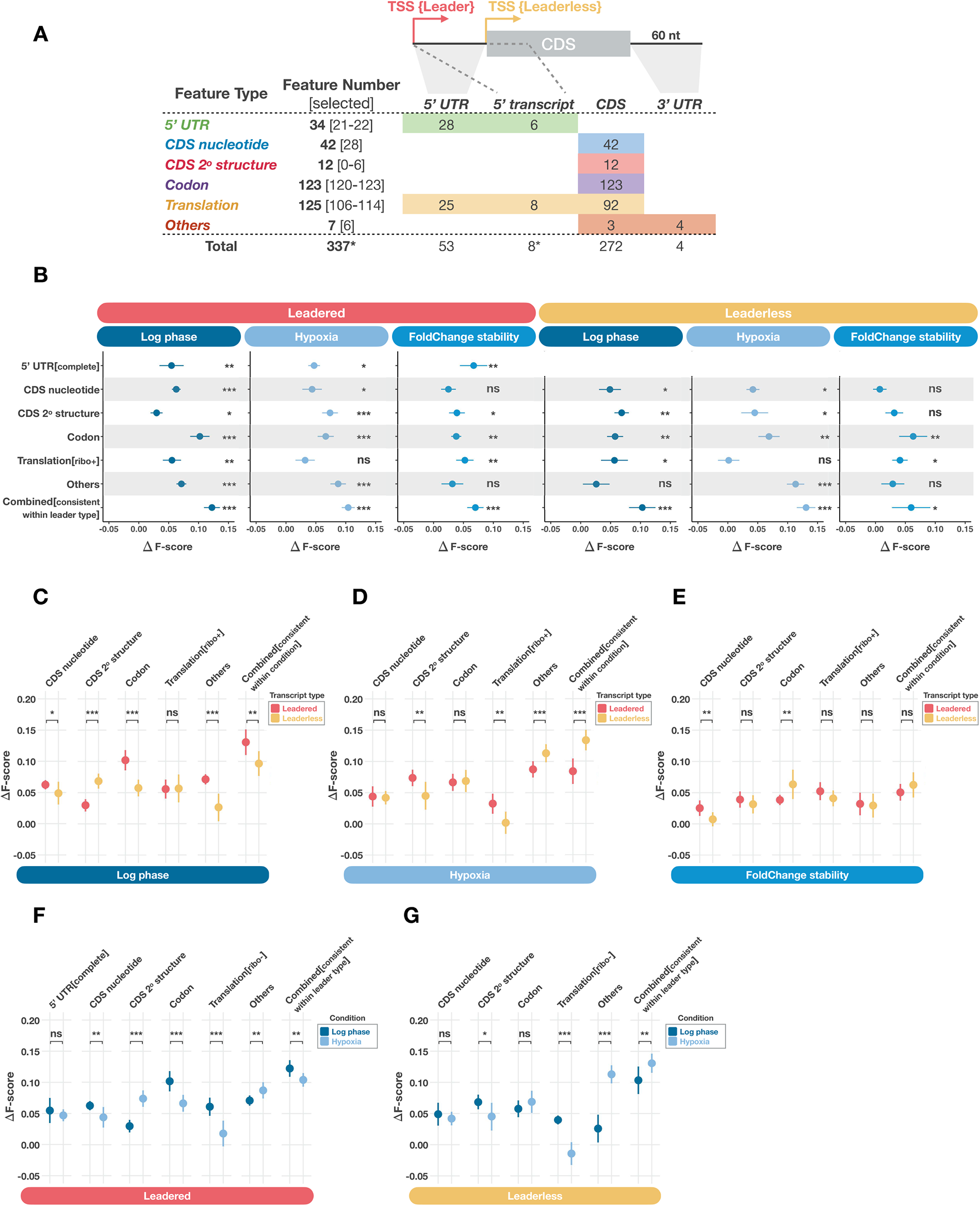
Non-linear combinations of diverse transcript properties specify half-life in *M. smegmatis*. **A.** Summary of transcript features used for random forest classifiers. The features were grouped into six types and quantified for specific transcript regions. Numbers in square brackets indicate the number of features of each type selected by our feature selection process. Numbers in the colored boxes indicate the number of features of each type in each transcript region. Asterisks indicate cases where the total number of unique features is less than the sum of the numbers above because some features are classified as 5’ UTR-type features for leadered transcripts and translation-type features for leaderless transcripts (see Table S3). **B.** Comparisons of classifier performance to random prediction models. Random forest classifiers were trained separately for leadered and leaderless transcripts to predict stability class in three conditions using various feature sets. The combined feature sets were selected by the log phase model for each transcript type (see Materials and Methods) and were used to train models in all three conditions. The 5’ UTR feature set includes both translation-related and non-translation-related features. The translation feature set includes log phase ribosome profiling. ΔF-score represents the difference in averaged F-score between random forest classifiers and random prediction estimators. Dots and bars represent mean and standard deviation of ΔF-scores for 10 repetitions of each model. The significance of the performance differences between random forest classifiers and random prediction estimators was tested using Nadeau and Bengio’s corrected paired t-test (Materials and Methods). *p* < 0.05 *, *p* < 0.01 **, *p* < 0.001 ***. **C-E.** Comparisons of ΔF-scores between leadered and leaderless transcript random forest models in log phase, hypoxia, and fold change in hypoxia relative to log phase. For each condition, the combined feature sets were selected by the leaderless model and were used to train models of both transcript types. **F-G.** Comparisons of ΔF-scores between log phase and hypoxia random forest models for leadered and leaderless transcripts. For each transcript type, the combined feature sets were selected by the log phase model and were used to train models of both log phase and hypoxia. The translation feature set excludes log phase ribosome profiling. The significance of the differences in model performance in **C-G** were tested using the Wilcoxon rank-sum test (Materials and Methods). For all panels, *p* < 0.05 *, *p* < 0.01 **, *p* < 0.001 ***.

Our results confirmed the association of 5’ UTR features with transcript stability as suggested in multiple studies ^8,12,21,22,25,27–35^. We also found that translation-related features were more important in log phase than in hypoxia for both transcript types, which is further explored below. Notably, ΔF-scores resulting from the combined feature sets were far less than the sum of the ΔF-scores from the individual feature types, indicating that the collective effect was not a result of linearly accumulated contributions of each feature type. This is consistent with the idea that transcript features interact in a non-linear fashion with respect to their impact on transcript half-life. Additionally, and in contrast to some previous reports ^2,21^, we found that no individual feature or feature type appeared to be a dominant determinant of half-life. Rather, our results indicate that the underpinnings of mRNA stability in *M. smegmatis* are complex, arising from non-linear combinations of diverse transcript properties.

### The importance of secondary structure and translation in predicting mRNA half-life varies by transcript type and condition

To determine if some feature types were differentially important depending upon transcript type (leadered or leaderless) or condition, we directly compared the performances of classifiers for leadered vs leaderless transcripts in each condition (Figure 3C-E) and for log phase vs hypoxia for each transcript type (Figure 3F, G). To rigorously compare ΔF-scores, the same feature set should be used in the models being compared. However, there were cases where the features differed between models, such as the absence of 5’ UTR features in the leaderless gene models. We therefore compared the leadered and leaderless models to each other using classifiers trained with only the features that were present in both (Figure 3C-E).

When considering the feature types separately, we found that CDS secondary structure features (as measured by the ΔG of minimum free energy, “MFE,” structures, ΔGMFE) were significantly more important for leaderless transcripts than for leadered transcripts in log phase (Figure 3C), which was exactly the opposite of the situation in hypoxia (Figure 3D). In direct comparisons between conditions the same trend was observed for CDS secondary structure features, which were more important in hypoxia than log phase for the leadered genes but more important in log phase than in hypoxia for the leaderless genes (Figure 3F, G). These results indicate that secondary structure differentially contributes to the stability of leadered and leaderless transcripts in different conditions.

A different pattern was seen for codon features, which were more important for leadered genes than for leaderless genes in log phase only, and more important in log phase than in hypoxia for leadered genes only (Figure 3C, F). The impact of codon content on half-life is likely related at least in part to translation having a greater influence on half-life in log phase, as observed for both transcript types in Figure 3B. However, comparing the impact of translation-related features between conditions was complicated by the inclusion of ribosome profiling data, which was performed only in log phase. We therefore trained classifiers excluding ribosome profiling features (Figure 3F, G) and directly compared their performance in log phase vs hypoxia for each transcript type. We still observed better performance of translation features in log phase than in hypoxia, suggesting that translation has a larger impact on transcript half-life in log phase than in hypoxia regardless of the transcript type.

We hypothesized that translation influenced transcript half-life more in log phase because in that condition most mRNAs were being translated at rates that varied according to transcript properties, while in hypoxia most transcripts were not being actively translated. For technical reasons, we tested this experimentally using carbon starvation rather than hypoxia. In previous work, we found that carbon starvation induced transcript stabilization similar to that seen in hypoxia ^39^. Here we performed polysome profiling and found that indeed, while monosomes and polysomes were readily detected in log phase cells, they had much lower abundance relative to ribosomal subunits in carbon-starved cells (Supplementary Figure S5). We furthermore collected fractions from the polysome profiling gradients and used qPCR to compare the relative abundance of four arbitrarily selected mRNAs in various fractions. For all four transcripts, the amount of transcript associated with monosomes and polysomes compared to unbound transcript decreased in carbon starvation compared to log phase (Supplementary Figure S5). These results are consistent with the idea that in non-growing cells, a larger portion of transcripts are unassociated with ribosomes compared to in actively growing cells.

To compare the collective effect of all features between leadered and leaderless transcripts, we trained classifiers for both using the same set of features selected for leaderless models in each condition (Figure 3C-E). The combined features were better able to predict stability for leadered transcripts than for leaderless transcripts in log phase, and vice versa in hypoxia (Figure 3C, D). Similarly, we compared the collective effect of all features between log phase and hypoxia using the same sets of features selected for log phase models for each transcript type (Figure 3F, G). The result showed better performance in log phase than hypoxia for leadered transcripts and the opposite for leaderless transcripts, consistent with the results of the direct transcript type comparisons. For the classifiers predicting the extent of stabilization in response to hypoxia, we observed no significant difference between leadered and leaderless transcripts for the majority of the shared feature types except CDS nucleotide and codon content features (Figure 3E). However, the leadered/leaderless comparison by necessity excluded 5’ UTR features, and we noted that the 5’ UTR features were the feature type that best predicted fold change stability for leadered genes (Figure 3B). This contrasted with the individual log phase and hypoxia classifiers where the 5’ UTR feature group was relatively weak (Figure 3B). Overall, our results indicate that the specific ways that various properties contribute to transcript stability are tied to the leader type as well as the condition.

### Identification of specific features differentially predictive of half-life for leadered and leaderless transcripts

In order to identify the transcript features that were differentially important for classification of leadered vs. leaderless transcripts, we evaluated the Gini importance rankings of the same set of features, selected by leaderless models, when used to train both leadered and leaderless models. For each condition, we combined the top 20 most important features identified in the leadered and leaderless models and compared the relative importance levels of these features for the two gene types (Log phase, Figure 4A; Hypoxia and fold-change in hypoxia, Supplementary Figure S6A, B; Compare to 20 least important features in log phase, Supplementary Figure S6C). We found that the most important features included features from each of the feature types in all three conditions, which further confirmed the collective effect of many features on dictating transcript stability. Furthermore, these comparisons also highlighted the different importance levels of many of the features between transcript types.

**Figure 4.**
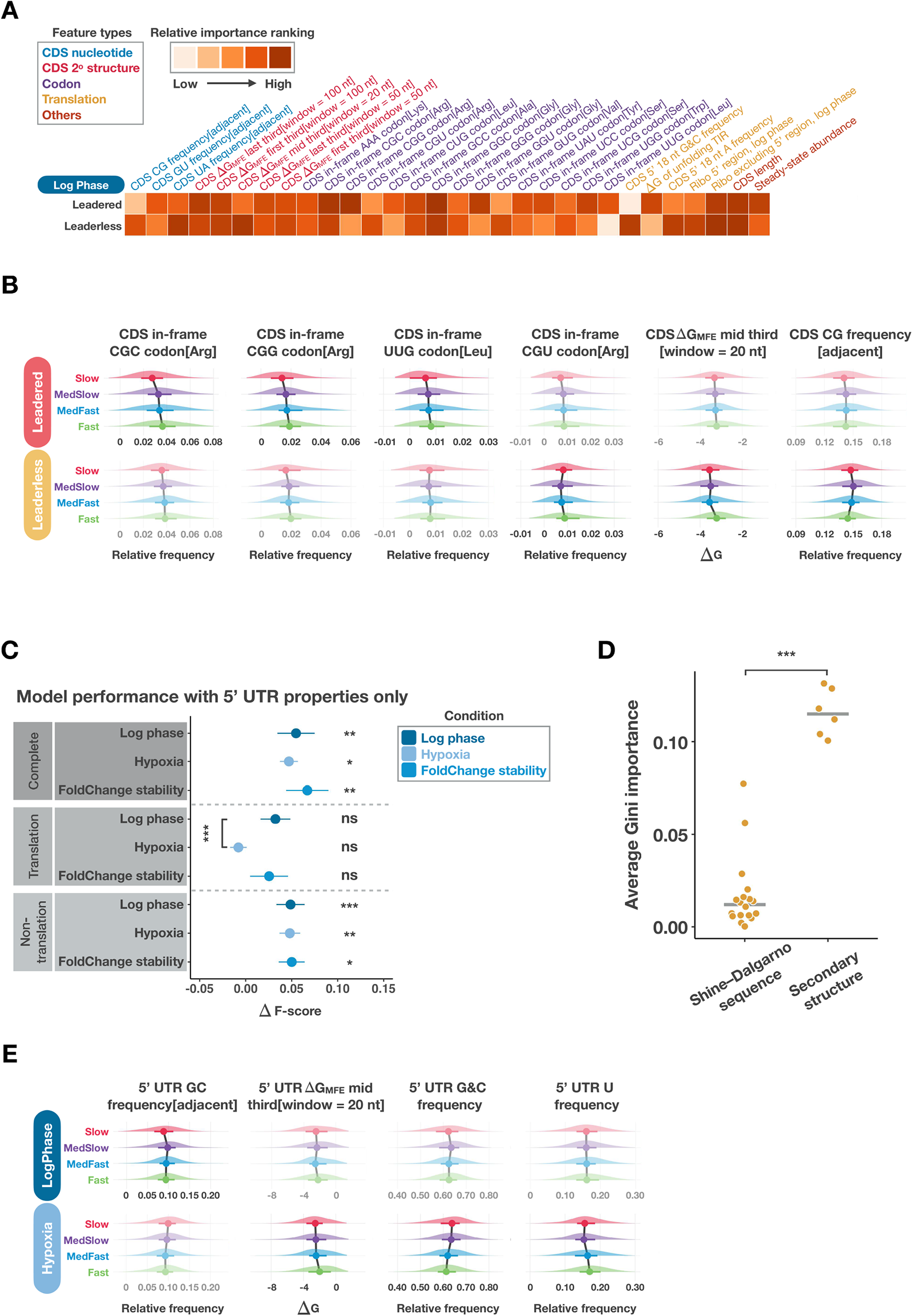
Transcript features differentially predict half-life for leadered and leaderless transcripts in log phase. **A.** Summary of the most important features for the leadered and leaderless half-life class prediction models in log phase. Random forest classifiers were trained using the same set of features, selected by the leaderless model, for leadered and leaderless transcripts. The 20 features with the highest Gini importance rankings for each model were combined and their relative importance rankings indicated by intensity of coloration in the heatmap. See Table S3 for feature definitions and details. **B.** Feature value distributions within each half-life class for selected features that differentially predicted half-life class for leadered and leaderless transcripts. Dimmed plots indicate that the feature was less important for that gene type. Dots and bars represent median and interquartile range. **C.** Comparisons of leadered gene models using only 5’ UTR features in three conditions. Models were trained and compared using the complete set of 5’ UTR features, translation-related 5’ UTR features only, or non-translation-related features only. See Table S3 for the specific features in each category. The performance differences between random forest classifiers and random prediction estimators were tested using Nadeau and Bengio’s corrected paired t-test. The log phase and hypoxia models using translation-related features were compared to each other with the Wilcoxon rank-sum test. **D.** Comparison of the importance of Shine-Dalgarno sequence features and secondary structure features in the ribosome binding regions of 5’ UTRs in the log phase model for leadered transcripts. Each dot is the average Gini importance value from 10 repetitions of the model. The difference in Gini importance was tested using the Wilcoxon rank-sum test. **E.** Examples of 5’ UTR features that differentially predicted half-life class between log phase and hypoxia. For all panels, *p* < 0.05 *, *p* < 0.01 **, *p* < 0.001 ***.

To determine the specific relationships between features of interest and half-life, we plotted the feature value distributions for each stability class (Figure 4B. SHAP distributions in Supplementary Figure S10). This allowed us to better understand why these features were important for model predictions and, more interestingly, how they were associated with transcript stability. Consistent with our finding that codon frequencies were more important for leadered than leaderless transcripts in log phase, we observed a number of specific codons with higher importance levels for leadered compared to leaderless transcripts. Among them, CGC (Arg), CGG (Arg) and UUG (Leu) are examples of codons with higher importance for leadered transcripts. Their distributions exhibited inverse correlations with stability for both leadered and leaderless transcripts, suggesting that they may negatively impact transcript stability (Figure 4B). However, these inverse relationships were stronger for leadered transcripts than for leaderless transcripts, which may explain the differences in importance for the classifiers (Figure 4B). In contrast, another Arg codon, CGU, was more important for leaderless transcripts compared to leadered transcripts and had a more complex relationship with half-life class (Figure 4B).

For leaderless transcripts, both the frequency of CG dinucleotide motifs and extent of CDS secondary structure were positively correlated with stability (Figure 4B). These trends were weaker for leadered genes. These results support the conclusion that in log phase, CDS secondary structure plays a more important role for leaderless transcripts compared to leadered transcripts.

### 5’ UTRs appear to influence transcript half-life through both translation-related and translation-independent mechanisms

The differences in stability determinants between leadered and leaderless transcripts were not limited to these shared features. Although we showed that the 5’ UTR itself was capable of predicting transcript stability (Figure 3B, Supplementary Figure S6D), the mechanisms by which it impacts stability were unclear. To further explore this, we categorized 5’ UTR features as translation-related (e.g., Shine-Dalgarno sequence and predicted secondary structure in the ribosome binding region) and non-translation-related (e.g., nucleotide content and predicted secondary structure outside of the ribosome binding region) and trained models separately using these two feature groups (Figure 4C, Supplementary Figure S6E). Surprisingly, our results suggest that the non-translation-related features have a larger impact on transcript stability than the translation-related features in both log phase and hypoxia. However, the model performance of translation-related features was significantly better in log phase compared to hypoxia, which is consistent with our finding that translation is more important for predicting transcript stability in log phase. Among all the translation-related features in 5’ UTR, the secondary structure seemed to be more important than the Shine-Dalgarno sequence (Figure 4D), which is the opposite of what was previously reported in *E. coli* ^21^. Such a difference could be because of the GC-richness of mycobacteria, which may cause secondary structure to have a bigger impact on ribosome access compared to less GC-rich species.

Consistent with our finding that CDS secondary structure was more important in hypoxia for leadered transcripts, several 5’ UTR features associated with secondary structures were more predictive of transcript half-life in hypoxia. The overall 5’ UTR G+C frequency was positively correlated with stability, while 5’ UTR ΔGMFE and U nucleotide frequency were negatively correlated with stability (Figure 4E, SHAP distributions in Supplementary Figure S10). The frequency of the GC dinucleotide motif showed a similar trend although it was more predictive in log phase. For both the GC dinucleotide and the overall G+C content, the relationships with half-life were monotonic in hypoxia but were more complex in log phase, with the slowest half-life class having lower frequencies than the medium-slow class. This could be a result of GC-rich sequences producing secondary structure that reduces ribosome binding in some cases. Given the greater apparent impact of translation on half-life in log phase, we expect that impediments to ribosome binding would negatively affect half-life in log phase more than in hypoxia.

### Leaderless gene start codons appear to affect transcription rate but not transcript half-life

Mycobacteria use both AUG and GUG start codons at high frequency. However, leaderless transcripts have more GUG start codons while leadered transcripts have more AUG start codons (Supplementary Figure S7A, D), leading us to investigate the relationship between start codon and half-life. Start codon identity had low Gini importance rankings for both leadered and leaderless genes, suggesting that it may not be a major determinant of translation efficiency for either transcript type. Despite its low Gini importance, AUG-initiating leadered transcripts had slightly longer half-lives on average than GUG-initiated leadered transcripts in log phase (Supplementary Figure S7B, E). When we examined steady-state transcript abundance as a function of start codon, we found that GUG-initiated transcripts had higher abundance on average, and that this effect was substantially stronger for leaderless transcripts (Supplementary Figure S7C, F). Since the relationship between start codon usage and steady-state abundance was stronger for leaderless transcripts and not explained by half-life, we considered that the identity of the first nt of a transcript may affect the efficiency of transcription initiation. It is well known that *E. coli* RNA polymerase preferentially initiates transcription with purines ^64^, and consistent with this, mycobacterial transcripts most often begin with purines ^5,6,38^. We examined the identity of the first nt of the 5’ UTRs of leadered transcripts and found that while transcripts beginning with As and Gs had equivalent half-lives, those beginning with G had higher average abundance (Supplementary Figure S7G-I). Together, these data suggest that mycobacterial RNA polymerase initiates transcription more efficiently with GTP than ATP.

### Identification of specific features differentially predictive of half-life in log phase and hypoxia

In order to identify the specific transcript features that were differentially important for classification among conditions, for each transcript type we used the same set of features, selected by log phase models, was used to train models in log phase, hypoxia and fold change in hypoxia. We then combined the top 20 features from each condition and compared the relative importance levels of these features across conditions (Figure 5A). Our results further confirmed the collective effect of various features on dictating transcript stability in each condition, but more importantly, revealed the ways in which the contributions of these features differed among conditions. Consistent with results of training with 5’ UTR related features only (Figure 3B, 4C), the 5’ UTR features remained important across conditions in the combined feature models for leadered transcripts (Figure 5A). These results further support the idea that 5’ UTRs influence transcript stability.

**Figure 5.**
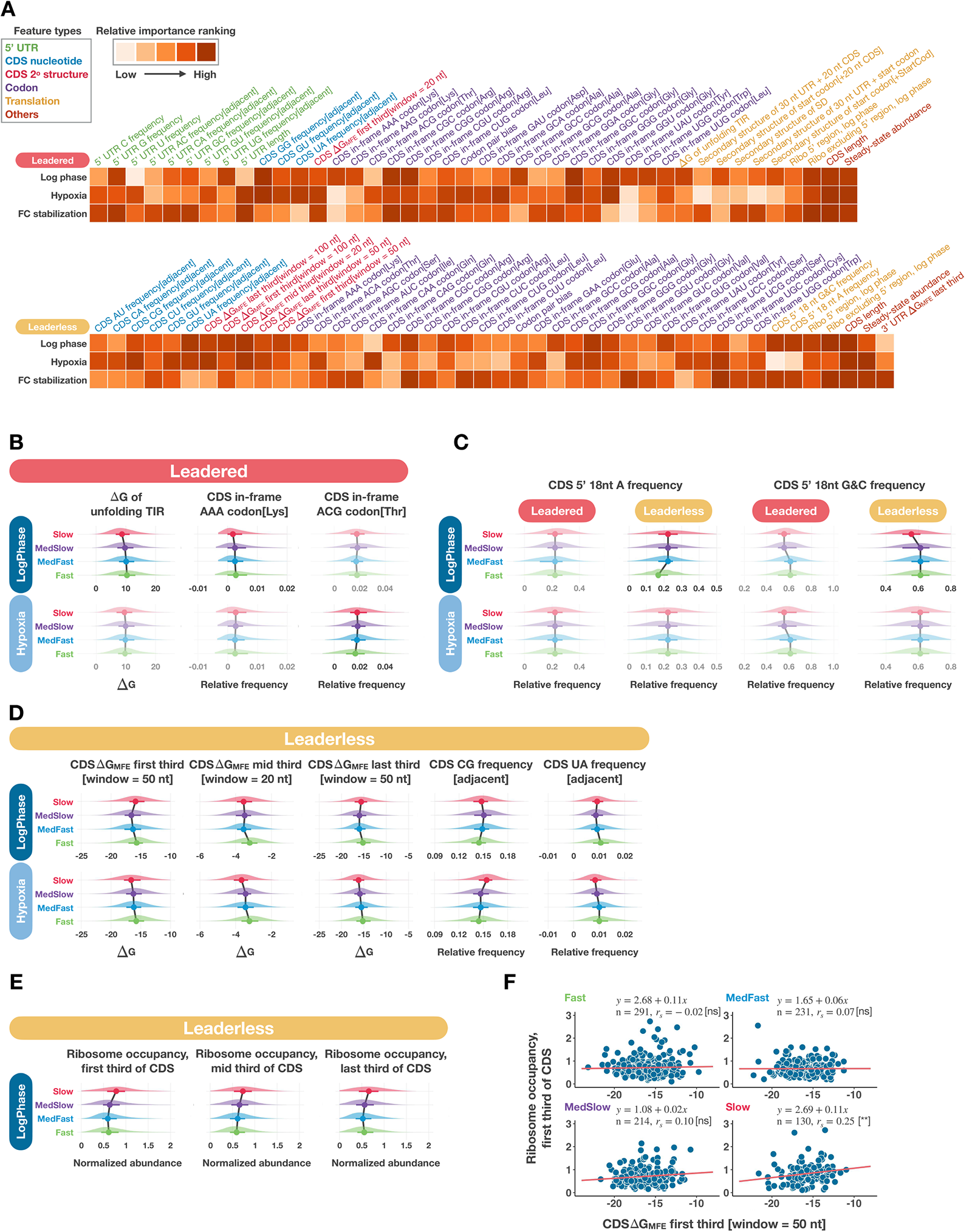
Transcript features differentially predict half-life in log phase and hypoxia. **A.** Summary of the most important features for the log phase, hypoxia, and log-to-hypoxia-fold-change models for leadered and leaderless transcripts. For each transcript type, random forest classifiers were trained for all three conditions using the set of features selected by the log phase model. For each transcript type, the 20 features with the highest Gini importance scores in each condition were then combined and their relative importance rankings indicated by intensity of coloration in the heatmap. See Table S3 for feature definitions and details. **B-D.** Feature value distributions within each half-life class for selected features that differentially predicted half-life class in different models. Dimmed plots indicate that the feature was less important for that condition and/or gene type. Dots and bars represent median and interquartile range. **B.** Selected features that were differentially important for log phase and hypoxia models for leadered genes. **C.** Selected features that were more important for log phase leaderless transcript models and are expected to impact the secondary structure of the 5’ ends of coding sequences. Plots for leadered genes are shown for comparison even though these features were not highly ranked for any leadered transcript models. **D.** Selected secondary-structure-related features that were relatively highly ranked for leaderless genes in both log phase and hypoxia models but showed different patterns of distributions across half-life classes for the two conditions. **E.** Log phase ribosome occupancy was quantified separately for each third of the CDS of each leaderless transcript. The x axes denote abundance of reads from ribosome-bound RNAs mapping to the indicated transcript regions. **F.** For leaderless genes, the log phase ribosome occupancy for the first third of each CDS was plotted as a function of the ΔGMFE of the first third of the CDS. rs denotes Spearman correlation, with the statistical significance in square brackets. *p* < 0.01 **, *p* < 0.001 ***.

The ΔG of unfolding secondary structure at translation initiation regions (TIRs) is a feature that can be used to predict the ribosome accessibility ^10^. We found that the ΔG of unfolding TIRs was an important transcript feature associated with half-life for leadered transcripts in log phase (Figure 4A, 5B, SHAP distributions in Supplementary Figure S11). In line with our finding that translation was more important in log phase, the ΔG of unfolding TIRs exhibited a stronger inverse correlation with half-life in log phase than in hypoxia (Figure 5B). This is consistent with a model in which higher accessibility of TIRs to ribosomes leads to greater translation efficiency or greater association of transcripts with ribosomes, thus protecting transcripts from degradation in log phase. The greater importance of translation in log phase was also supported by the stronger correlation between frequencies of certain codons and half-life, such as AAA (Lys) (Figure 5B). However, the effects of codon frequency on transcript half-life might be a mixture of translational and non-translational effects, as suggested by the higher importance of ACG (Thr) in hypoxia (Figure 5B), where translation overall appears to have less impact on half-life.

In contrast, for leaderless transcripts in log phase, our results indicated a more complicated relationship between secondary structure and translation, and their correlations with transcript stability. Although it was not reflected by the ΔG of unfolding TIRs, 5’ end secondary structure was important for leaderless transcripts in log phase. Particularly, we observed a low A nucleotide frequency in the first 18 nucleotides of the CDS for transcripts in the fast half-life class and a low G+C frequency for those in the slow half-life class (Figure 5C, SHAP distributions in Supplementary Figure S11). Notably, these features associated with secondary structure of the first 18 nt of CDSs were more predictive of half-life class for leaderless than leadered genes. While low secondary structure in this region is typical in many organisms ^63^ and was experimentally shown to increase translation efficiency for leadered transcripts in *E. coli* ^62^, it may have a larger influence on translation of leaderless genes because these lack the additional ribosome recruitment signals found in 5’ UTRs.

We found that the impact of secondary structure continued beyond the 5’ 18 nt of leaderless transcripts. We calculated ΔGMFE with different sequence window sizes to measure the secondary structure of the 5’ third, middle third, and 3’ third of each CDS. Consistent with our previous observation that these features were collectively more predictive of half-life class for leaderless genes in log phase (Figure 3C), we found generally negative correlations between CDS ΔGMFE and transcript half-life (Figure 5D, SHAP distributions in Supplementary Figure S11). These correlations were maintained when ΔGMFE was calculated using different window sizes (Supplementary Figure S8A). The trends of CG dinucleotide and UA dinucleotide frequency also supported this idea (Figure 5D). Overall, our results suggested that the CDS secondary structure is positively correlated with transcript half-life regardless of the region of CDS, consistent with the idea that secondary structure generally protects transcripts from cleavage by RNases. These relationships were monotonic in hypoxia, but more complex in log phase where transcripts in the slow half-life class deviated from the otherwise monotonic trend, having less secondary structure than those in the medium-slow class (Figure 5D). We hypothesized that stronger secondary structure might compete with ribosome binding, and since translation appears to have a strong protective effect in log phase only, transcripts in the slow class in log phase might be protected more by ribosome binding than by secondary structure. To test this, we quantified the ribosome occupancy within the 5’ third, middle third, and 3’ third of each CDS and evaluated their correlations with transcript half-life. As expected, we observed that the slow half-life class had the highest average ribosome occupancy across the entire CDS (Figure 5E). The idea of competition between secondary structure and ribosome binding was further supported by a positive correlation between ΔGMFE and ribosome occupancy for the first third of transcripts in the slow class (Figure 5F). This trend was maintained when ΔGMFE was calculated using a different window size, but was not observed for the middle and 3’ thirds of transcripts (Supplementary Figure S8B-E). Together, our results highlight the complexity of interplay between transcript features.

### Transcript abundance and length are more predictive of half-life in hypoxia

Among the most important features, steady-state abundance and CDS length were identified by models across transcript types and conditions (Figure 4A, 5A, Supplementary Figure S6A, B). The relationship between transcript abundance and half-life has been investigated in various bacteria, yet the results are conflicting (Reviewed in ^65^). Broadly consistent with some studies in *M. tuberculosis* ^2^, *E. coli* ^20–22,66–68^ and *L. lactis* ^19,22,69^, we found that the distributions of transcript abundance exhibited an inverse correlation with half-life for both transcript types and conditions (Figure 6A, SHAP distributions in Supplementary Figure S12). However, the correlations we observed were substantially weaker than what was reported for *M. tuberculosis* in log phase ^2^. Interestingly, in *M. smegmatis* the inverse correlations between transcript abundance and half-life were substantially stronger in hypoxia than log phase for both transcript types (Figure 6B). Comparing the correlations for leadered and leaderless transcripts did not reveal differences in log phase, but a stronger correlation was seen for leaderless transcripts compared to leadered transcripts in hypoxia (Figure 6B). These results indicate that, underlying the broad inverse correlation between transcript abundance and half-life, the strength of the relationship varies depending on condition and to a lesser extent on transcript type.

**Figure 6.**
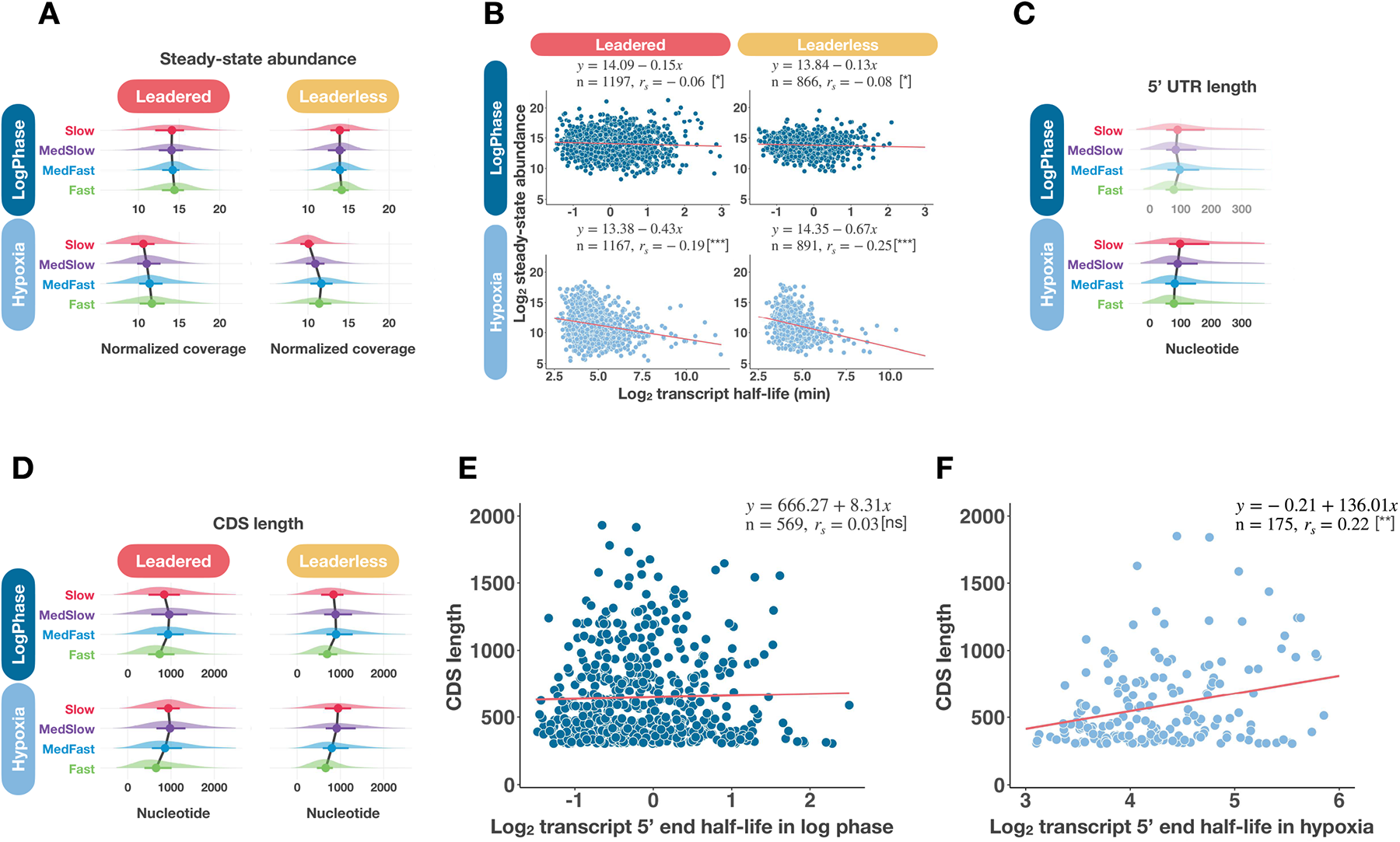
Steady-state transcript abundance is negatively correlated with half-life, while transcript length is positively correlated with mRNA half-life in hypoxia. **A.** Distributions of steady-state transcript abundance within each half-life class in log phase and hypoxia for leadered and leaderless transcripts. **B.** Correlations between steady-state abundance and transcript half-life. While abundance was highly ranked in all models (see Figure 5A), its negative correlation with half-life was stronger in hypoxia. **C.** Distributions of 5’ UTR lengths within half-life classes for leadered transcripts in log phase and hypoxia. This feature had a high importance ranking in hypoxia only. **D.** Distributions of CDS length within each half-life class in log phase and hypoxia. This feature was highly ranked in all models. **E-F.** Half-lives were calculated for only the first 300 nt of each CDS and genes were selected that had similar half-lives for the 5’ 300 nt and the whole CDS (see Figure S9). For these subsets of genes in log phase and hypoxia, the correlation between CDS length and 5’ 300 nt half-life are shown. In **B, E, F**, rs denotes Spearman correlation, with the statistical significance in square brackets. *p* < 0.05 *, *p* < 0.01 **, *p* < 0.001 ***.

5’ UTR length was an important feature in hypoxia but not in log phase (Figure 5A), and consistent with this, had a clear monotonic positive correlation with transcript half-life in hypoxia (Figure 6C, SHAP distributions in Supplementary Figure S12). In contrast, the relationship between 5’ UTR length and transcript half-life was weaker and less straightforward in log phase (Figure 6C). CDS length was an important feature for both transcript types in both conditions (Figure 5A), exhibiting a roughly monotonic positive relationship with half-life class in hypoxia but a non-monotonic relationship in log phase (Figure 6D, SHAP distributions in Supplementary Figure S12). This is consistent with the finding in *M. tuberculosis* that CDS length has little broad correlation with transcript half-life in log phase ^2^. In contrast, CDS length has also been shown to have negative correlation with transcript half-life in *L. lactis*, *E. coli*, and *S. cerevisiae* ^19,21,24^. While the strong predictive power of CDS length in *M. smegmatis* was intriguing, there were two potential confounding factors. First, RIF only inhibits promoter escape following transcription initiation, having no impact on elongating RNA polymerases. We attempted to control for this during the process of half-life determination by identifying transcripts with delays in degradation following the addition of RIF and excluding the 1-2 minute delay periods from the half-life calculation (see Figure S2 in ^38^). However, for longer transcripts the elongating RNA polymerases may continue for longer than 2 minutes, leading to an overestimation of half-life. Secondly, recent studies of transcript 3’ ends in *M. tuberculosis* ^70^ and *E. coli* ^71^ suggested that a sizable fraction of transcripts present in cells are degradation intermediates or incomplete transcripts resulting from premature transcription termination or paused RNA polymerases. We cannot distinguish these from complete transcripts in our RNAseq libraries, and it is possible that longer transcripts give rise to more incomplete transcripts that are long enough to be captured in RNAseq libraries and these have different degradation kinetics than complete transcripts.

To account for these potential confounders, we calculated the half-life of only the first 300 nt of each CDS (the “5’ end half-life”). For each condition, we then divided genes into five groups according to the ratio of the 5’ end half-life to the entire gene half-life (“log2 half-life ratio”, Supplementary Figure S9A, D). We also calculated the steady-state (0 minute RIF) RNAseq coverage ratio of the 5’ 300 nt versus 3’ end 300 nt of each gene within these groups (Supplementary Figure S9A, D). As expected, those genes with differential abundance of transcript 5’ and 3’ regions often had non-zero log2 half-life ratios, consistent with the idea that incomplete transcript fragments often have different degradation kinetics than full-length transcripts. On the other hand, genes with similar coverage of their 5’ and 3’ 300 nt generally had similar log2 half-life ratios (Supplementary Figure S9A, D, groups 3 and 4 respectively), indicating that these genes are likely less affected by the confounders described above (Colored group in Supplementary Figure S9A, D). For these non-confounded genes, there was no correlation between 5’ end half-life and CDS length in log phase (Figure 6E), but there was a significant positive correlation in hypoxia (Figure 6F). These relationships were maintained when leadered and leaderless transcripts were analyzed separately (Supplementary Figure S9B, C, E, F). Consistent with the idea that the positive correlation in hypoxia was due to the condition rather than the selection of genes, we found little to no correlation between 5’ end half-life and CDS length for the genes in Figure 6F in log phase (Supplementary Figure S9G-I). In summary, although able to contribute to model predictions for both transcript types in log phase, transcript abundance and CDS length seemed to have stronger correlations with half-life in hypoxia. Consistent with this, the 5’ UTR length exhibited a positive correlation with half-life in hypoxia, suggesting that the overall transcript length is more important for predicting half-life in hypoxia than in log phase.

## DISCUSSION

In this study, we used transcriptome-wide mRNA half-life datasets to investigate the intrinsic features that impact transcript stability in aerobically growing and hypoxia-arrested *M. smegmatis*. This led us to discover the diverse transcript features that were differentially associated with mRNA stability depending on the microenvironment. Our results indicate that translation likely has a larger impact on mRNA degradation in log phase than in hypoxia. We further found that, coupled with the impact of conditions, transcript leader type (leadered vs leaderless) also impacted transcript stability through various transcript features. Importantly, our results showed that it is the collective effect of diverse transcript features that shapes the transcript stability landscape of *M. smegmatis*, with no single feature dominating. A collective impact of transcript features on mRNA half-life has been reported in other organisms as well ^21,24,25^, but in some studies transcription rate (as inferred from steady-state abundance and half-life) appeared to be a dominant feature with a much greater impact than other features ^2,21^.

Additionally, our study further revealed the non-linear character of interactions between features and ways in which their impacts differ according to transcript type and growth condition.

We developed machine learning models with the goal of associating transcript features with half-life by predicting half-life using a wide-ranging feature set, as well as to further identify likely determinants of transcript half-life by quantifying the strength of their associations.

Initially, we attempted to develop regression models given the continuous nature of transcript half-life values. However, the models failed to provide accurate prediction of half-life values as needed to draw reliable conclusions about feature relationships with transcript half-life.

Therefore, we grouped transcript half-life values into four classes to predict half-life through classification instead. The decision to define four half-life classes was informed by the hierarchical clustering of degradation profiles to estimate the number of groups to best represent transcriptome-wide stability. Besides the innate difficulty of the four-class prediction task, the intertwined non-linear correlations, existing not only between transcript features and half-life, but also among transcript features themselves, make the classifications even more challenging. Despite the difficulties, our models achieved significantly better performance than random predictions. To compensate for the suboptimal model performance, we implemented SHAP visualization to enhance our interpretation of model predictions by showing the prediction direction along with the feature values for each half-life class (Supplementary Figure S10-S12). We found that these results were consistent with the correlations observed from the distributions of individual transcript features. Together, these computational tools provided us with enough confidence and information to draw conclusions about the associations between transcript features and half-life. Nonetheless, our current models still lack the ability to fully explain the relationships between transcript features themselves, and the mechanism of how they work together to determine transcript stability.

Future studies on the relationships among those important transcript features will greatly improve model predictions and advance our understanding of the mechanisms regulating transcript degradation.

Like *M. tuberculosis*, *M. smegmatis* exhibited variance in transcript half-lives during log phase growth and showed transcriptome-wide stabilization when exposed to hypoxia (Figure 2B) ^2,39^. Despite the potential differences in regulatory mechanisms between species, our study of *M. smegmatis* still provides insights to facilitate understanding of transcript stability in *M. tuberculosis*. Unlike in the previous study in *M. tuberculosis* ^2^, we were able to quantify mRNA half-lives transcriptome-wide in hypoxia, showing that the extent of stabilization varied among genes and indicating that the determinants of half-life differ between the two conditions. It was reported that transcript abundance was the single feature strongly correlated with transcript half-lives in log phase in *M. tuberculosis*, while features like CDS length and G+C content showed little correlation ^2^. Here, we greatly expanded the scope of candidate features and found diverse transcript features that could contribute to predicting half-life in *M. smegmatis*. Whether the collective and differential effect of the wide range of transcript features on half-life we observed in *M. smegmatis* also exists in *M. tuberculosis* awaits further investigation.

Our results also suggested that the lack of broad correlations between transcript features and half-lives could be due to not only condition, but also to transcript-type-dependent regulatory mechanisms in mycobacteria.

In log phase, transcripts in both *M. tuberculosis* and *M. smegmatis* exhibited little correlation between CDS length and half-life. However, we found that the correlation became stronger in hypoxia for *M. smegmatis*. A previous study suggested that motion of large cytoplasmic components was dramatically reduced in *Caulobacter crescentus* and *E. coli* when metabolic activity was reduced, due to decreased fluidity of the cytoplasm ^72^. The positive correlation between CDS length and half-life is consistent with this idea, as longer transcripts would be more affected by the reported changes in diffusion rates ^72^, leading to reduced encounters between transcripts and RNases in the hypoxic cytoplasm.

Transcript abundance was another feature whose influence was affected by growth condition. Similar to *M. tuberculosis* ^2^, we observed an inverse correlation between transcript abundance and half-life in log phase, although this relationship was much weaker in *M. smegmatis*. However, we found that the strength of the correlation was stronger in hypoxia compared to log phase. Such an association has been reported for other bacteria as well ^19–22,66–69,73,74^, although conflicting reports exist for *E. coli*, where some report a negative correlation ^20–22,66–68^ while others report no correlation or a positive correlation ^73,75^. The mechanistic basis of the negative correlation reported in many studies is unknown, although it has been suggested to be a function of the impact of transcript abundance on encounters with RNases ^21,22^.

It has been shown in *E. coli* and *S. cerevisiae* that translation efficiency is positively correlated with mRNA half-life in log phase ^76,77^. Our results also provided evidence to support this association in *M. smegmatis* as we observed translation-related features were more important for half-life predictions in log phase compared to hypoxia. Besides the previously identified differences in translation mechanisms between leadered and leaderless transcripts ^10,11,78^, our results indicate that mRNA degradation mechanisms may also differ between leadered and leaderless transcripts. We first confirmed that 5’ UTR features were predictive of mRNA half-life in leadered transcripts. There were also differences in the importance of CDS features in predicting half-lives of leadered vs leaderless transcripts. For example, G+C content was particularly low in the first 18 nt of the CDS specifically for leaderless transcripts with the slowest half-lives, consistent with the idea that secondary structure in this region has a larger impact on translation efficiency for leaderless transcripts compared to leadered transcripts.

In summary, our results suggest that underlying the observed transcript stability patterns in mycobacteria lies a complex interplay between inherent transcript features and microenvironments. Additionally, our study provides a foundation to facilitate further investigation of transcript stability in mycobacteria, as well as an experimental and computational framework to study transcript stability more broadly in other organisms.

## DATA AVAILABILITY

All RNAseq data generated in this study are available at GSE227248. Other data and code generated for analysis in this study are available from the following GitHub repository, https://github.com/ssshell/mRNA_stability.

## AUTHOR CONTRIBUTIONS

H.S., Y.Z., D.A.V.-B., D.K., and S.S.S designed the experiments and computational analysis. Y.Z. and D.A.V.-B. performed experiments and generated data. H.S., C.S.M., J.M.K., and J.K.M. conducted computational analyses. D.K. and S.S.S supervised the research and interpreted the results. H.S., D.K., and S.S.S wrote the manuscript.

## ACKNOWLEDGEMENTS

We thank members of the Shell and Korkin labs for helpful discussions.

