## Supplementary methods for "Diverse intrinsic properties shape transcript stability and stabilization in *Mycolicibacterium smegmatis*"

### SUPPLEMENTAL METHODS

#### Culture conditions for polysome profiling

For carbon starvation cultures, cells were grown to log phase ($OD_{600}$ = 0.8) in rich medium, pelleted, and rinsed three times with carbon starvation medium (Middlebrook 7H9 with 5 g/L BSA, 0.85 g/L NaCl, 3 mg/L catalase, and 0.05% Tyloxapol) at 4°C and then resuspended in carbon starvation medium to an $OD_{600}$ of 0.8 and incubated at 200 rpm and 37°C. *M. smegmatis* remained in these conditions for 22 hours before been used for RNA determination experiments. Log phase control cultures were grown in the same medium with the addition of 2 g/L glucose and 0.2% glycerol, and Tween-80 in place of Tyloxapol.

#### Polysome profiling

Biological triplicate cultures of *M. smegmatis* in logarithmic phase ($OD_{600}$ = 0.94, 300 mL) or in 22-h carbon starvation ($OD_{600}$ = 0.94, 600 mL) were filtered over 0.20 µm filters (VWR part number 10040-468) using a vacuum pump at room temperature (~23°C), and cells were scraped into 50 mL of liquid nitrogen. Lysis buffer (20 mM Tris pH 8, 10 mM $MgCl_2$, 100 mM $NH_4Cl$, 5 mM $CaCl_2$, 0.4% Triton-X 100, 0.1% Igepal (NP-40), 1 mM chloramphenicol, and 100 U/mL RNase-free DNase, in DEPC water) was carefully added to the same liquid nitrogen, forming small crystal spheres. Frozen cell pellets and crystalized lysis buffer were ground in a Retsch CryoMill using 10 mL grinding jars and a single 7 mm stainless steel ball (6 cycles of 3 min at 15 Hz, with a 1 min pause in between). Cell lysates were thawed at 30° for 2 min and then on ice for 30 min. Lysates were clarified twice by centrifugation at 21,130 x g and 4°C for 10 min. 5%-50% linear sucrose gradients in 100 mM NaCl, 10 mM $MgCl_2$, and 30 mM Tris-HCl pH7.5 in DEPC water were made using a Biocomp Gradient Master. Lysate containing 1 mg of RNA diluted in RNase-free water to a final volume of 150 µL was layered onto sucrose gradients in triplicate. Sucrose gradients were centrifuged in an Optima L90K ultra centrifuge (SW 41 Ti rotor) at 35,000 RPM (151,000 x g) for 2 h 45 min at 4°C, and analyzed using a Gradient Profiler (Biocomp) with an EM-1 Econ UV Monitor (Bio-Rad) while collecting 150 µL fractions.

**Quantitative PCR (qPCR) of polysome profiling fractions**

Normalized transcript degradation A spike-in RNA was made by in vitro transcription and added to polysome profiling fractions prior to RNA extraction for qPCR. The *mCherry* (M10) sequence + T7 terminator (782 bp) was obtained from plasmid pSS374 using EcoRI. RNA was *in vitro* generated with a HiScribe T7 Quick High Yield RNA Synthesis Kit (NEB, E2050S) followed by RNA purification using the LiCl protocol, according to the manufacturer's instructions. 1 ng of this *mCherry* mRNA was added to 80 µL of each polysome profiling fraction, RNA was purified using acid phenol:chloroform:IAA (125:24:1, pH 4-5), and cDNA was synthesized as described in[1]. qPCR was performed as described in[1] using the primers described therein for *sigA*, *atpE*, and *rnj*. Other primer sets were Ms1: GCCGGAAGAGAAGGCTAGAT and CGTCCGCTTTTCGAAACTAC; 16S: AAGCGCAAGTGACGGTATGTG and AAGCTGTGAGTTTTCACGAACAAC; 23S: AGCCTGTAGGGAGTCAGATAG and GCAGCATAGGATCACCGAAT; *mCherry*: GATGGTGTAGTCCTCGTTGTG and GAGGTCAAGACCACCTACA. Abundance for all other transcripts was calculated relative to *mCherry*.

**Hierarchical clustering to identify transcript degradation patterns**

Normalized transcript degradation profiles in log phase and hypoxia were collected as described above and in [2] for clustering analysis. To select genes with high quality degradation profiles, we conducted two preprocessing procedures. For each gene, we first calculated the coefficient of variation (CV) over three RNAseq replicates for each time point. Genes with CV > 0.75 in any of the first 4 time points (0, 1, 2, 4 minutes for log phase; 0, 3, 6, 9 minutes for hypoxia) were excluded from clustering. In order to cluster by the differences in degradation pattern rather than the absolute abundance, we then converted the profiles into relative abundance to an initial time point (0 and 1 minute for log phase; 0 minute for hypoxia). The CV-filtered genes were further selected by being required to have relative abundance at subsequent timepoints be no more than 1.5 times that of the initial time point. After preprocessing, the relative degradation profiles were then clustered using hierarchical clustering with the Euclidean distance measure and ward.D2 agglomeration method. The degradation pattern of each cluster was represented by the mean and standard deviation of $log_2$ mRNA abundance at each timepoint (Supplementary Figure S2B-D and S2F-H). mRNA degradation is expected to follow a single exponential decay trend. For log phase, initial clustering produced a cluster of genes that exhibited a delay prior to the start of exponential decay, which is a well-established phenomenon due to rifampicin blocking transcription initiation but not elongation [3]. To produce clusters unaffected by this technical issue, we removed the genes in the cluster showing the delay and re-clustered the remaining genes using degradation profiles normalized to the 1 minute timepoint rather than the 0 minute timepoint. This resulted in classification of 4972 genes into degradation pattern classes in log phase (Supplementary Figure S2A). For hypoxia, no delays were observed, likely due to both transcription and mRNA degradation being substantially slower than in log phase, and classes were directly defined for 5098 genes by the clustering of degradation profiles relative to the 0 minute timepoint (Supplementary Figure S2E).

### Feature selection algorithm

Listed below is a pseudocode of our algorithm to reduce the number of correlated features.

**Algorithm:** Feature selection for a given feature set $\boldsymbol{F_f}$ with class labels

**Input:** Complete feature set $\boldsymbol{F_f}$

**Metrics:**

[$\rho_{ff}$], Spearman's rank correlation coefficient to quantify correlation between the values of each pair of features.

[$\tau_{fc}$], Kendall rank correlation coefficient to quantify correlation between the values of each feature and the class. To calculate Kendall rank correlation, class labels are converted to numerical representations (*e.g.* "Fast" to 1, "Med-fast" to 2, "Med-slow" to 3, "Slow" to 4).

[$\boldsymbol{Ave_{adjustedP}}$], Mean FDR adjusted *P* value of Kruskal-Wallis test (KW) and Kolmogorov-Smirnov test (KS). The KW test compares the values of a given feature among classes. The KS test adjusted *P* value is taken as the minimum FDR adjusted *P* value of all pair-wise comparisons of values for a given feature between classes.

[$\boldsymbol{N_{corr}}$], number of features correlated with a given feature.

[$\boldsymbol{N_{sigTest}}$], number of statistically significant tests (KW, KS) of a given feature in comparisons among/between classes.

**Procedure:**

1. Preprocessing. Removed features with zero variance.
2. Get correlated feature sets $\boldsymbol{F_{corr}}$. Each $\boldsymbol{F_{corr}}$ included features with | $\rho_{ff}$ | >= 0.6. Only $\boldsymbol{F_{corr}}$ that meet following criteria are further considered for selection:
   a. Correlated features have the same transcript region [*5′ UTR/5′ transcript, CDS, 3′ UTR*] and the same feature type [*Nucleotide, Codon, Secondary structure, Ribosome, Others*]
3. Determine the order in which $\boldsymbol{F_{corr}}$s will be evaluated. For all the $\boldsymbol{F_{corr}}$ that meet the criteria in step 2, first evaluate the $\boldsymbol{F_{corr}}$ that includes the feature that has the highest | $\tau_{fc}$ | among all features in all $\boldsymbol{F_{corr}}$. If tied, start with the $\boldsymbol{F_{corr}}$ that has the least amount of correlated features.
4. Evaluate and select the best feature from each $\boldsymbol{F_{corr}}$. For features within $\boldsymbol{F_{corr}}$:
   a. Select the feature that meets criteria in the following order:
      i. Maximum $\boldsymbol{N_{sigTest}}$. If tied, go to the next metric,
      ii. Minimum $\boldsymbol{N_{corr}}$. If tied, go to the next metric,
      iii. Minimum group sum | $\rho_{ff}$ |. If tied, go to the next metric,
      iv. Maximum | $\tau_{fc}$ |. If tied, go to the next metric,
      v. Minimum $\boldsymbol{Ave_{adjustedP}}$
   b. Update the rest of $\boldsymbol{F_{corr}}$ by removing the features that were not selected.
5. Continue until all $\boldsymbol{F_{corr}}$ have been evaluated.

**Output:** the list of selected features.

**<u>Machine learning classifier training algorithm</u>**

Listed below is a pseudocode of a generalized classifier training algorithm. For our classifiers, $\boldsymbol{k}$ = 5, $\boldsymbol{n}$ = 10.

**Algorithm:** Classifiers training and evaluation with ***k***-fold nested cross-validation for ***n*** repetitions.

**Input:** Feature set ***D*** with class labels, Hyperparameter set ***H***

1. For ***i = 1 … n*** repetitions:
2. Training(***D***, ***k***, ***H***):
3. Random stratified partition ***D*** into ***k*** folds $\boldsymbol{D_1 \dots D_k}$
4. For ***j* = 1 … *k*** folds:
5. *TrainSet* = $\boldsymbol{D \setminus D_j}$
6. *TestSet* = $\boldsymbol{D_j}$
7. Train the RandomBaselineModel on *TrainSet*
8. ***h**** = RandomizedSearchCV[*TrainSet*, ***H***, RandomForestModel]
9. ***RandomForestModel**** = RandomForestModel trains on *TrainSet* using ***h****
10. $\boldsymbol{Fscore_j}$ **=** RandomBaselineModel & ***RandomForestModel**** predict on *TestSet*
11. $\boldsymbol{FeatureImportance_j}$ = Gini importance[***RandomForestModel****]
12. $\boldsymbol{SHAP_j}$ = SHAP values of ***RandomForestModel**** predicts on *TestSet*
13. $\boldsymbol{Fscore\_allFold_i}$ = [$\boldsymbol{Fscore_j}$]
14. $\boldsymbol{Fscore_i}$ = Mean[$\boldsymbol{Fscore_j}$]
15. $\boldsymbol{Gini_i}$ = Mean[$\boldsymbol{FeatureImportance_j}$]
16. $\boldsymbol{SHAP_i}$ = Concatenate[$\boldsymbol{SHAP_j}$]
17. ***Fscore_allFold*** = [$\boldsymbol{Fscore\_allFold_i}$]
18. ***Fscore_average*** = Mean[$\boldsymbol{Fscore_i}$]
19. ***Gini*** = Mean[$\boldsymbol{Gini_i}$]
20. ***SHAP*** = Concatenate[$\boldsymbol{SHAP_i}$]
21. Return ***Fscore_allFold***, ***Fscore_average***, ***Gini***, ***SHAP***, ***Metrics_others***

**Output:** Classifier performance of all folds of all repetitions ***Fscore_allFold (n = 50)***, classifier performance averaged across folds for each repetition ***Fscore_average (n = 10)***, impurity-based feature importance quantification averaged across repetitions ***Gini***, SHAP values of individual classes ***SHAP***. Output also includes other performance metrics such as precision and recall values for both individual class and averaged value across folds.
