## Supplementary material for "Diverse intrinsic properties shape transcript stability and stabilization in *Mycolicibacterium smegmatis*": Table S3

### *M. smegmatis feature table*

| Feature | | Feature type | Feature region | Feature column id | Numbers of feature | TL (transcript leader) status for genes have the feature | Comments |
|---|---|---|---|---|---|---|---|
| **Nucleotide** | percentage of nucleotide usage | Translation | CDS | Nucl_***nucleotide***_***region*** | 4 | Leadered, Leaderless, Leader_undefined | usage percent of each nucleotide of the ***5' end 18 nts of CDS*** |
| | | CDS Nucleotide | | | 8 | | usage percent of each nucleotide of the ***entire CDS (including last 3 nts), 3' end 18 nts of CDS (including last 3 nts)*** |
| | | 5' UTR | 5' UTR | Nucl_***nucleotide***_5pUTR | 4 | Leadered | usage percent of each nucleotide of the 5' UTR |
| | percentage of adjacent nucleotide usage | Translation | CDS | adja_***dinucleotide***_***region*** | 16 | Leadered, Leaderless, Leader_undefined | usage percent of each possible adjacent nucleotide combination of ***5' end 18 nts of CDS*** |
| | | CDS Nucleotide | | | 32 | | usage percent of each possible adjacent nucleotide combination of ***entire CDS (including last 3 nts), 3' end 18 nts of CDS (including last 3 nts)*** |
| | | 5' UTR | 5' UTR | adja_***dinucleotide***_5pUTR | 16 | Leadered | usage percent of each possible adjacent nucleotide combination of the 5' UTR |
| | nt G+C percentage | Translation | CDS | GC_***region*** | 1 | Leadered, Leaderless, Leader_undefined | GC usage of ***5' end 18 nts of CDS*** |
| | | CDS Nucleotide | | | 2 | | GC usage of ***entire CDS (including last 3 nts), 3' end 18 nts of CDS (including last 3 nts)*** |
| | | 5' UTR | 5' UTR | GC_5pUTR | 1 | Leadered | GC usage of 5' UTR |
| **Codon** | percentage of nonstop codons usage in CDS | Translation | CDS | ***codon***_***region*** | 61 | Leadered, Leaderless, Leader_undefined | usage percent of each nonstop codon of the ***5' end 18 nts of CDS*** |
| | | Codon | | | 122 | | usage percent of each nonstop codon of the ***entire CDS (excluding stop codon), 3' end 18 nts of CDS (excluding stop codon)*** |
| | start codon | Translation | CDS | start_codon_***codon*** | 3 | | binary indicator of start codon usage (***AUG, GUG, UUG***) |
| | stop codon | Translation | CDS | stop_codon_***codon*** | 3 | | binary indicator of stop codon usage (***UAA, UAG, UGA***) |
| | codon pair bias | Codon | CDS | CodonPairBias | 1 | | codon pair bias calculated using codon pair score of each codon pair in CDS (Coleman JR *et al. Science*. 2008) |
| **Secondary Structure** | MFE | Others | 3' UTR | tprUTR_MFE_20_10nt_***region*** | 4 | Leadered, Leaderless, Leader_undefined | 3'UTR is defined as 60 nts after stop codon. Calculate the average minimum free energy of subsequences from ***entire sequence, 5' end (~1/3), middle (~1/3) and 3' end (~1/3) of sequence***.The entire sequence is sliced into 20 nts windows, each time with 10 nts overlap. See calculating MFE for CDS. |
| | | 5' UTR | 5' transcript | transcript_5p20nt_MFE | 1 | Leadered | calculate minimum free energy of 5' end 20 nts of the transcripts (5' UTR for Leadered, CDS for Leaderless or 5' UTR plus CDS if UTR is shorter than 20 nts) |
| | | Translation | | | | Leaderless | |
| | | CDS MFE | CDS | CDS_MFE_***M***_***N***nt_***region*** | 12 | Leadered, Leaderless, Leader_undefined | calculate the average minimum free energy of subsequences from ***entire CDS, 5' end (~1/3), middle (~1/3) and 3' end (~1/3) of CDS***. Slicing CDS into ***M*** nts (***20, 50, 100***) windows, each time with ***N*** nts (***10, 25, 50***) overlap. Take the average of all the subsequences' MFE as the final number. If the remaining subsequence is shorter than window size, then use the whole remaining subsequence to get MFE without further slicing. For partial CDSs (~1/3), take the average of the subsequence windows that starting within the partial region. (***e.g.*** For 102 nts CDS, 5' end is [1, 34], middle is [35, 68], 3' end is [69, 102]. [***50***, ***25***] slicing windows are {[1, 50], [26, 75], [51, 100], [76, 102]}. The 5' end will take the average MFE of {[1, 50], [26, 75]}, middle will be the MFE of [51, 100], 3' end will be the MFE of [76, 102]). In cases that there's no subsequence window starting within the middle or 3' end region, it will take the last available subsequence window from upstream region for the current region. |
| | | 5' UTR | 5' UTR | fprUTR_MFE_20_10nt_***region*** | 4 | Leadered | For 5' UTR longer than 35 nts, extract 5' UTR sequence excluding the last 15 nts. Calculate the average minimum free energy of subsequences from ***entire sequence, 5' end (~1/3), middle (~1/3) and 3' end (~1/3) of sequence***.The entire sequence is sliced into 20 nts windows, each time with 10 nts overlap. See calculating MFE for CDS. |
| | number of unpaired nt at 5' end | Translation | CDS | CDS_5pUnpairedNt | 1 | Leadered, Leaderless, Leader_undefined | number of unpaired base at 5' end of the sequence when folding entire CDS, 5' UTR, 5' UTR plus 18 nts of 5' end of CDS, 5' end 20 nts of the transcripts (5' UTR for Leadered, CDS for Leaderless or 5' UTR plus CDS if UTR is shorter than 20 nts) |
| | | 5' UTR | 5' UTR | fprUTR_5pUnpairedNt | 1 | Leadered | |
| | | | | fprUTR_plus18ntCDS_5pUnpairedNt | 1 | Leadered | |
| | | 5' UTR | 5' transcript | transcript_5p20nt_5pUnpairedNt | 1 | Leadered | |
| | | Translation | | | | Leaderless | |
| | probability of unpairing | Translation | 5' UTR | ***seq***_***method***_***region***_UnpairedProb | 6 | Leadered | for leadered genes with 5' UTR longer than 30 nts, the unpaired probability of ***SD (-6 to -14), entire folding region*** (***both are average for the whole sequence of the probabilities that each nucleotide and the nucleotide before it are both unpaired***), ***start codon*** (***the probability that whole start codon unpaired***) by folding ***last 30 nts of 5' UTR plus first 20 nts of 5' end of CDS & last 30 nts of 5' UTR plus start codon*** |
| | | 5' UTR | 5' transcript | transcript_5p20nt_***method***_***region***_UnpairedProb | 4 | Leadered | the unpaired probability of ***first 3 nts, first 5 nts*** of 5' end (probability of 3 nts or 5 nts are ***all unpaired & the average of the probability that each nt is unpaired***) by folding 5' end 20 nts of the transcripts (5' UTR for leadered, CDS for leaderless or 5' UTR plus CDS if UTR is shorter than 20 nts) |
| | | Translation | | | | Leaderless | |
| | MFE unfold of translation initiation region | Translation | 5' transcript | MFE_unfold | 1 | Leadered, Leaderless | calculate $\Delta G_{unfold}$ ($\Delta G_{init}$ - $\Delta G_{mRNA}$) of translation initiation regions (TIRs) for Leadered (5' UTR >= 12 nts) and Leaderless (Bharmal MM *et al. NAR Genom Bioinform*. 2021) |
| **Ribosome** | ribosome occupancy | Translation | 5' transcript | ribo_***region*** | 1 | Leadered, Leaderless, Leader_undefined | ribosome profiles (raw TPM ratio (Totalribo/TotalmRNA), NOT log2 scale, normalized by total mRNA; YX Chen *et al. Proc Natl Acad Sci.* 2020) of ***CDS, CDS plus 20 nts upstream of 5' end, 5' end of transcripts*** (5' end 18 nts of Leaderless, 20 nts upstream plus 5' end 18 nts of Leadered and Others), ***CDS excluding 5' end 18 nts*** |
| | | Translation | CDS | | 3 | | |
| | Shine-Dalgarno GA features | Translation | 5' UTR | SD_GA_***feature*** | 2 | Leadered, Leader_undefined | ***GA percent & GA count*** of SD region (-17 to -4 relative to start codon). GA percent is the percentage of Gs and As in the sequence. GA count is the total frequency of all di-nucleotide sequences (GG, AA, AG, and GA) in two "reading frames" of the sequence, one starting at the first nt and one starting at the second nt of the sequence. (***e.g.*** For sequence 'AGGATCTCTAAAGG', first frame 'AG GA TC TC TA AA GG' has 4 G+A di-nts, second frame 'GG AT CT CT AA AG' has 3 G+A di-nts. The GA count for this sequence would be 7) |
| | Shine-Dalgarno motif frequency | Translation | 5' UTR | ***motif***_5p25ntUpStream | 17 | | motif frequency (***'AGAAAGGAGGT', 'AGGA', 'AGGAG', 'AGGAGG', 'GAAAGG', 'GAGG', 'GGAG', 'GGAGG', 'AAGGAG', 'AAGGA', 'AAAGGA', 'AAGG', 'AAAGG', 'GAAAG', 'GAAA', 'AGGAA', 'GGAA'***) of 25 nts upstream of the start codon |
| **Other Properties** | sequence length | Others | CDS | CDS_length | 1 | Leadered, Leaderless, Leader_undefined | length of CDS, 5' UTR (length is 0 for Leaderless [it is not used for training models of Leaderless], NA for Leader_undefined) |
| | | 5' UTR | 5' UTR | fpr_UTR_length | 1 | Leadered, *Leaderless* | |
| | initial abundance | Others | CDS | init_abund_***condition*** | 2 | Leadered, Leaderless, Leader_undefined | initial abundance (raw normalized nt sum coverage of $T_0$ [normalized by ***library size, qPCR normalization factors, CDS length***], NOT log2 scale, average over replicates) of ***log phase*** (con_noAtc) and ***hypoxia*** |
